# The Genomic History of the Middle East

**DOI:** 10.1101/2020.10.18.342816

**Authors:** Mohamed A. Almarri, Marc Haber, Reem A. Lootah, Pille Hallast, Saeed Al Turki, Hilary C. Martin, Yali Xue, Chris Tyler-Smith

## Abstract

The Middle East is an important region to understand human evolution and migrations, but is underrepresented in genetic studies. We generated and analysed 137 high-coverage physically-phased genome sequences from eight Middle Eastern populations using linked-read sequencing. We found no genetic traces of early expansions out-of-Africa in present-day populations, but find Arabians have elevated Basal Eurasian ancestry that dilutes their Neanderthal ancestry. A divergence in population size within the region starts before the Neolithic, when Levantines expanded while Arabians maintained small populations that could have derived ancestry from local epipaleolithic hunter-gatherers. All populations suffered a bottleneck overlapping documented aridification events, while regional migrations increased genetic structure, and may have contributed to the spread of the Semitic languages. We identify new variants that show evidence of selection, some dating from the onset of the desert climate in the region. Our results thus provide detailed insights into the genomic and selective histories of the Middle East.

## Introduction

Global whole-genome sequencing projects have provided insights into human diversity, dispersals, and past admixture events (Bergström *et al.*, 2020; Mallick *et al.*, 2016; GenomeAsia100K Consortium, 2019; 1000 Genomes Project Consortium *et al.*, 2015). However, many populations remain understudied, which restricts our understanding of genetic variation and population history, and may exacerbate health inequalities (Sirugo *et al.*, 2019). A region particularly understudied by large-scale sequencing projects is the Middle East. Situated between Africa, Europe and South Asia, it forms an important region to understand human evolution, history and migrations. The demographic history and prehistoric population movements of Middle Easterners are poorly understood, as are their relationships among themselves and to other global populations. The region contains some of the earliest evidence of modern humans outside Africa, with fossils dated to ~180 thousand years ago (kya) and ~85 kya identified in the Levant and North West Arabia, respectively (Hershkovitz *et al.*, 2018; Groucutt *et al.*, 2018). In addition, tool kits suggesting their presence have been identified in South East Arabia dating to ~125 kya (Armitage *et al.*, 2011). Although most of Arabia is a hyper-arid desert today, there were several humid periods resulting in a ‘green Arabia’ in the past which facilitated human dispersals, with the onset of the current desert climate thought to have started around 6 kya (Petraglia *et al.*, 2020). The toggling from humid to arid periods has been proposed to result in population movements adapting to the climate. The Neolithic transition within Arabia may have developed independently within the region, or resulted from an expansion of Levantine Neolithic farmers southwards (Drechsler, 2009; Uerpmann, *et al.,* 2010; Crassard *et al.*, 2013a; Crassard *et al.,* 2013b; Hilbert *et al.,* 2015). To address such questions, we generated and analysed a high-coverage physically-phased open-access dataset of populations from the Arabian Peninsula, the Levant and Iraq. In addition to creating a catalogue of genetic variation in an understudied region that will assist future medical studies, we have investigated the population structure, demographic and selective histories, and admixture events with modern and archaic humans.

## Results

### Dataset and Sample Sequencing

We sequenced 137 whole genomes from eight Middle Eastern populations (Figure 1A) to an average coverage of 32x using a library preparation method that preserves long-range information from short reads, and aligned them to the GRCh38 reference (Methods). An advantage of using this ‘linked-read’ technology is the reconstruction of physically-phased haplotypes and improved alignments at repetitive regions which confound short-read aligners (Figure S1). All populations investigated speak Arabic, a Semitic language of the Afro-Asiatic language family, with the exception of the Iraqi Kurdish group who speak Kurdish, an Iranian language belonging to the Indo-European family. After quality control (Methods) we identify 23.1 million single nucleotide variants (SNVs). We compared our dataset to variants identified in the recently released Human Genome Diversity Project (HGDP-CEPH) study (Bergström *et al.*, 2020). We find 4.9 million autosomal SNVs in our dataset that are not found in the HGDP. As expected, most of the new variants are rare (93%, < 1% minor allele frequency); however, ~370,000 are common (> 1%). Interestingly, most of these common variants are outside the accessibility mask defined by Bergstrom *et al*., 2020 (~246,000). This illustrates the importance of sequencing genetically under-represented populations such as Middle Easterners and the inclusion of regional-private variants in future medical studies. It also demonstrates that a significant amount of unknown variation resides in regions that are not accessible to standard short-reads.

**Figure 1.**
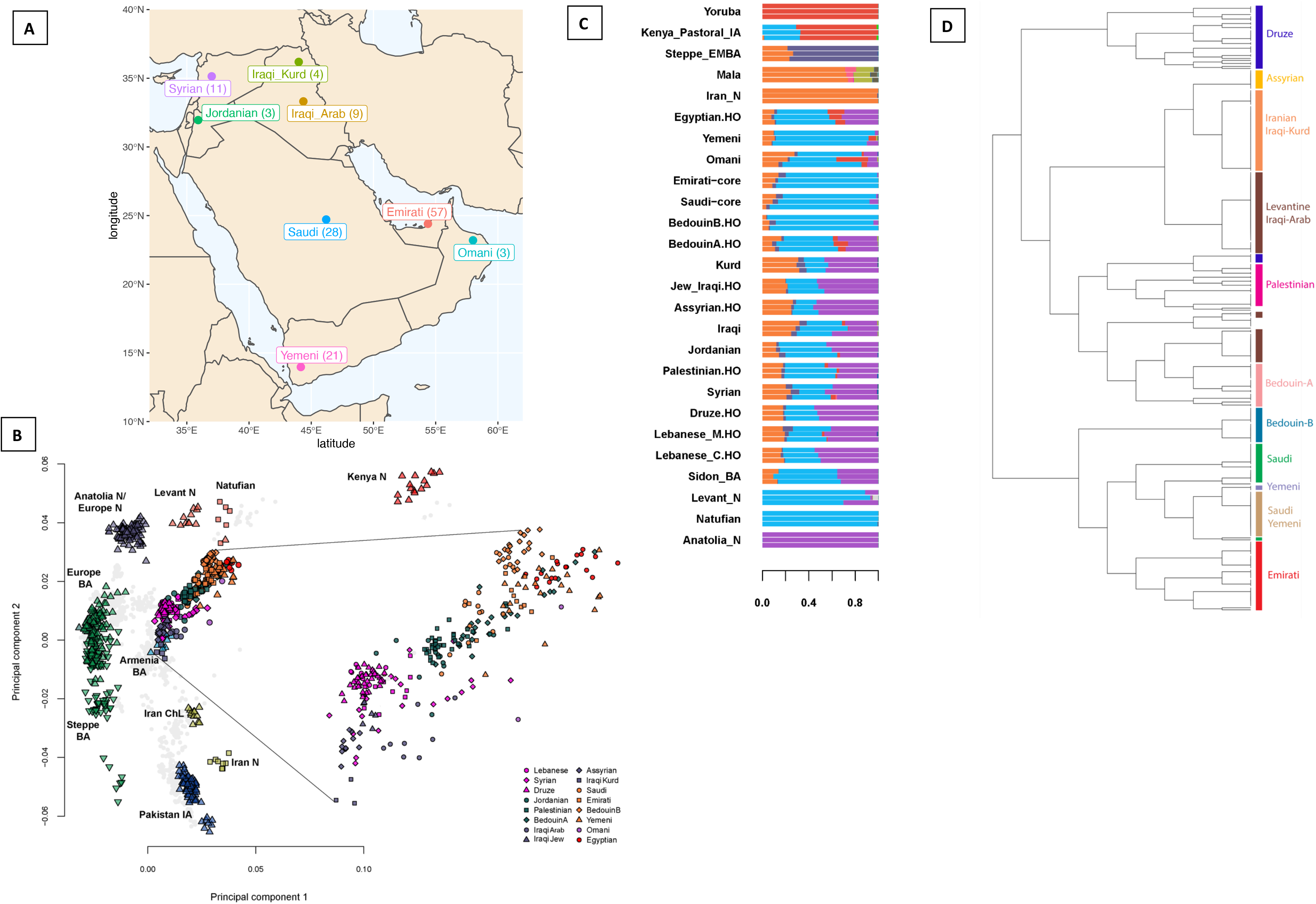
Overview of the dataset and population structure of the Middle East. **A)** Map illustrating the populations sampled in this study, with numbers in brackets illustrating number of individuals. **B)** Principal component analysis of ancient and modern populations. Eigenvectors were inferred with present-day populations from the Middle East, North and East Africa, Europe, Central and South Asia. The ancient samples were then projected onto the plot (all modern non-Middle Easterners shown as grey points). Plot also shows a magnification of the modern Middle Eastern cluster. See Figure S3 for more details. **C)** Temporally-aware model-based clustering using ∼80,000 transversions and 9 time points. Showing K=13 when the Anatolia_N and Natufian components split. See Figure S5 for more details. “.HO” suffix refers to samples from the Human Origins Dataset. **D)** Finestructure tree of modern-day Middle Easterners with population clusters highlighted. See Figure S2 for more details.

### Population Structure and Admixture

Uncovering population structure and past admixture events is important for understanding population history and for designing and interpreting medical studies. We explored the structure and diversity of our dataset using both single-variant and haplotype-based methods. After merging our dataset with global populations, fineSTRUCTURE (Lawson *et al.*, 2012) identified genetic clusters that are concordant with geography, and showed that self-labelled populations generally formed distinct clusters (Figure 1D and S2). Populations from the Levant and Iraq (Lebanese, Syrians, Jordanians, Druze and Iraqi-Arabs) clustered together, while Iraqi-Kurds clustered with Central Iranian populations. Arabian populations (Emiratis, Saudis, Yemenis and Omanis) clustered with Bedouins (BedouinB) from the HGDP. The fineSTRUCTURE analysis thus allowed us to identify subpopulations who show minimal admixture, which we herein label ‘core’.

We next analysed our samples in the context of ancient regional and global populations. Principal component analysis (Figures 1B and S3) shows that present-day Middle Easterners are positioned between ancient Levantine hunter-gatherers (Natufians), Neolithic Levantines (Levant_N), Bronze Age Europeans and ancient Iranians. Arabians and Bedouins are positioned close to ancient Levantines, while present-day Levantines are drawn towards Bronze Age Europeans. Iraqi Arabs, Iraqi Kurds and Assyrians appear relatively closer to ancient Iranians and are positioned near Bronze Age Armenians. We find that most present-day Middle Easterners can be modelled as deriving their ancestry from four ancient populations (Table S1): Levant_N, Neolithic Iranians (Iran_N), Eastern Hunter Gatherers (EHG), and a ~4,500 year old East African (Mota). We observe a contrast between the Levant and Arabia: Levantines have excess EHG ancestry (Figure S4), which we showed previously had arrived in the Levant after the Bronze Age along with people carrying ancient south-east European and Anatolian ancestry (Haber *et al.,* 2017, Haber *et al.* 2020). Our results here show this ancestry remained mostly confined to the Levant region. Another contrast between the Levant and Arabia is the excess of African ancestry in Arabian populations. We find that the closest source of African ancestry for most populations in our dataset is Bantu Speakers from Kenya, in addition to contributions from Nilo-Saharan speakers from Ethiopia specifically in the Saudi population. We estimate that African admixture in the Middle East occurred within the last 2,000 years, with most populations showing signals of admixture around 500-1,000 years ago (Figure S5 and Table S2).

In addition to differences in EHG and African ancestry, we observe an excess of Natufian ancestry in the South compared with the North (Figure S4). Model-based clustering also shows that Arabian populations have little Anatolia Neolithic (Anatolia_N) ancestry compared with the modern-day Levantines (purple component in Figure 1C). This result is intriguing since Levant_N shares significant ancestry with Anatolia_N compared with the preceding local Natufian population (Lazaridis *et al.*, 2016), and a hypothesized Neolithic expansion from the Levant to Arabia should have also carried Anatolia_N ancestry. The difference in ancient Anatolian ancestry could also be from post-Bronze Age events, which resulted in differences in EHG ancestry in the region (Haber *et al.*, 2020). When we substitute Levant_N with Natufians, we found that Arabians could be successfully modelled (Table S1 and Figure S7), suggesting that they could derive all of their local ancestry from Natufians without requiring additional ancestry from Levant_N. On the other hand, none of the present-day Levantines could be modelled as such.

In addition to the local ancestry from Epipaleolithic/Neolithic people, we find an ancestry related to ancient Iranians that is ubiquitous today in all Middle Easterners (orange component in Figure 1C; Table S1). Previous studies showed that this ancestry was not present in the Levant during the Neolithic period, but appears in the Bronze Age where ~50% of the local ancestry was replaced by a population carrying ancient Iran-related ancestry (Lazaridis *et al.*, 2016). We explored whether this ancestry penetrated both the Levant and Arabia at the same time, and found that admixture dates mostly followed a North to South cline, with the oldest admixture occurring in the Levant region between 3,900 and 5,600 ya (Table S3), followed by admixture in Egypt (2,900-4,700 ya), East Africa (2,200-3,300) and Arabia (2,000-3,800). These times overlap with the dates for the Bronze Age origin and spread of Semitic languages in the Middle East and East Africa estimated from lexical data (Kitchen *et al.*, 2009; Figure S8). This population potentially introduced the Y-chromosome haplogroup J1 into the region (Chiaroni *et al.*, 2010; Lazaridis *et al.*, 2016). The majority of the J1 haplogroup chromosomes in our dataset coalesce around ~5.6 [95% CI, 4.8-6.5] kya, agreeing with a potential Bronze Age expansion; however, we do find rarer earlier diverged lineages coalescing ~17 kya (Figure S9). The haplogroup common in Natufians, E1b1b, is also frequent in our dataset, with most lineages coalescing ~8.3 [7-9.7] kya, though we also find a rare deeply divergent Y-chromosome which coalesces 39 kya (Figure S9).

### Effective Population size and Separation History

Historical effective population sizes can be inferred through the distribution of coalescence times between chromosomes sampled from a population (Li and Durbin, 2011). However, there is limited resolution in recent periods using single human genomes, while errors in haplotype phasing create artefacts when using multiple genomes (Schiffels and Durbin, 2014; Terhorst *et al.*, 2017). Although methods have been developed that extend these approaches by incorporating the allele frequency spectrum from unphased genomes, they do not have resolution at recent times, for e.g. through the metal ages (Terhorst *et al.*, 2017; Bergström *et al.*, 2020). By leveraging recent advances in generating genome-wide genealogies (Speidel *et al.*, 2019), and the large number of physically-phased samples in our study, we could estimate the effective population size of each population in our dataset up to very recent times - 1 kya (Figure 2A and S16A). We found all Middle Easterners had a significant decrease in population size, around the out-of-Africa event ~50-70 kya. The recovery from this bottleneck follows a similar pattern until 15-20kya, when a contrast between the Levant and Arabia started to emerge. All Levantine and Iraqi populations continued to show a substantial population expansion, while Arabians maintained similar sizes. This contrast is noteworthy since it starts after the end of the Last Glacial Maximum and becomes prominent during the Neolithic, when agriculture developed in the Fertile Crescent and led to settled societies supporting larger populations. Following the Neolithic, and with the start of the aridification of Arabia around 6kya, Arabian populations experienced a bottleneck while Levantines continued to increase in size. The expansion in Levantines then plateaus and their population size decreases around the 4.2 kiloyear aridification event (Weiss et al., 1993). The decline in Emiratis is especially prominent, reaching an effective population size of ~5,000, more than 20 times smaller than Levantines and Iraqis at the same time period. A recovery can be observed in the past 2 ky.

**Figure 2.**
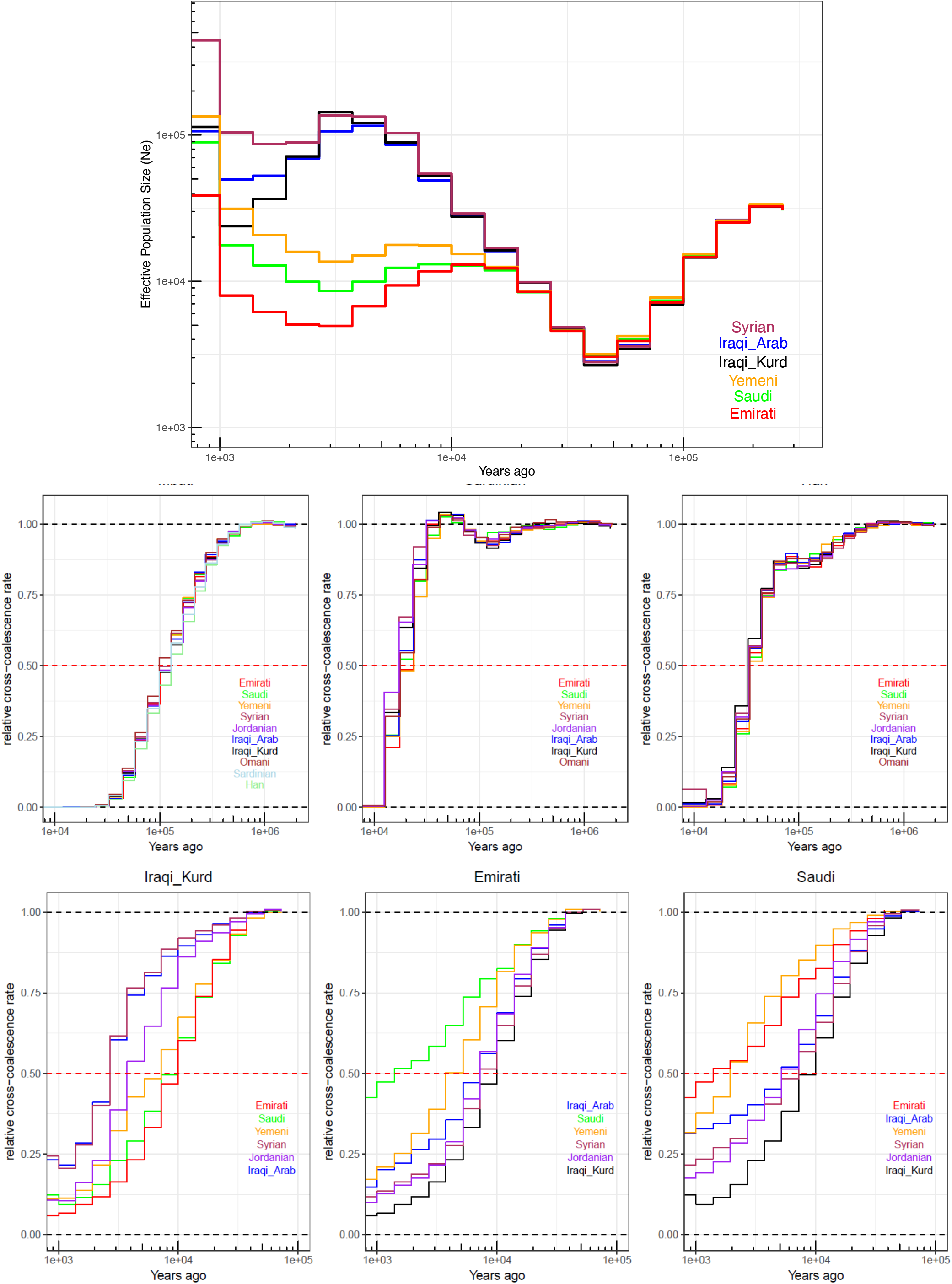
Population size and separation history. **Top)** Effective population size histories for Middle Eastern populations. More details in Figure S16A. **Center)** Separation history between Mbuti, Sardinians and Han (indicated at the top of each panel) with each of the Middle Eastern populations (identified within each panel). All Middle Eastern populations show similar split time with each of these global populations. **Bottom)** Separation history within the Middle East (population indicated at the top of each panel, and within each panel). More comparisons show in Figure S16B. Note the different X-axis scales.

We next studied the population separation history of Middle Eastern populations among themselves and from global populations. The importance of accurate phasing in this analysis is illustrated by an earlier finding that suggested, based on statistically phased data, that modern-day Papuans harbour ancestry of an early expansion of modern humans out of Africa (Pagani *et al.*, 2016). However, this was not replicated using physically-phased genomes, suggesting it was caused by a statistical phasing artefact (Bergström *et al.*, 2020). Conversely, when exploring population separation history at recent times, rare variants become more informative but are less accurately phased by statistical methods, and are unlikely to be present in reference panels. We first tested whether present-day Middle Easterners harbour ancestry from an early human expansion out of Africa by comparing the split times of our populations with physically-phased samples from the HGDP (Figure 2B and S10). Using a relative cross-coalescent rate (rCCR) of 0.5 as a heuristic estimate of split time, we found that Levantines, Arabians, Sardinians and Han Chinese share the same split time, and additionally the same gradual pattern of separation, from Mbuti ~120kya. We then compared the populations in our dataset with Sardinians and found they split ~20 kya, with Levantines showing a slightly more recent divergence than Arabians. In contrast to the gradual separation patterns to Mbuti, Sardinians show more of a clean split to all Middle Eastern populations. Notably, all lineages within the Levant and Arabia, and in addition to lineages within all Middle Eastern populations and Sardinians, coalesce within 40 kya. These results collectively suggest that present-day Middle Eastern populations do not harbour any significant traces from an earlier expansion out of Africa, and all descend from the same population that expanded out of the continent ~50-60 kya.

We then compared the separation times of populations within the Middle East, and found the oldest divergence times were between Arabia and the Levant/Iraq (Figure 2C and S16B). The Emiratis split from Iraqi Kurds around 10 kya, and more recently around 7 kya from Jordanians, Syrians and Iraqi Arabs. Saudi split times from the same populations appear more recent, around 5-7 kya, while the Yemeni separation curves are intermediate between the Emirati and Saudi curves. The split times between Arabia and the Levant predate the Bronze Age, agreeing with our phylogenetic modelling that if a Bronze Age expansion into Arabia occurred, it did not result in a complete replacement of ancestry.

Within the Levant and Iraq, all splits occurred in the past 3-4 ky. Within Arabia, Yemenis split from Emiratis ~4 kya and Saudis appear as the least divergent population to both the Emiratis and Yemenis, with recent splits within the last 2ky. The separation history of the region suggests continuous historical gene flow occurring between the Levant/Iraq and Central Arabia, and in addition between Central Arabia to the Southeast, and separately to the Southwest in Yemen.

### Archaic introgression and deep ancestry in the Middle East

The similar amount of Neanderthal ancestry in most non-African populations and the low diversity of introgressed haplotypes suggest that modern humans likely experienced a single pulse of Neanderthal admixture as they expanded out of Africa (Bergström *et al.*, 2020). Middle Eastern populations have previously been shown to have lower Neanderthal ancestry than European and East Asian populations (Rodriguez-Flores *et al.*, 2016; Bergström *et al.*, 2020); however, the interpretation of this finding is complicated by recent African admixture ‘diluting’ Neanderthal ancestry (Haber et al., 2016). In addition, some analyses require the use of an outgroup, which, if it itself contains Neanderthal ancestry, can bias estimates (Chen *et al.*, 2020). To investigate Neanderthal introgression in our dataset, we exploited the accurate phasing of our samples and compared cross-coalescent rates with the high coverage Vindija Neanderthal genome (Prüfer *et al.*, 2017). All Middle Easterners showed an archaic admixture signal at a time point similar to other Eurasians (Figure 3A).

**Figure 3.**
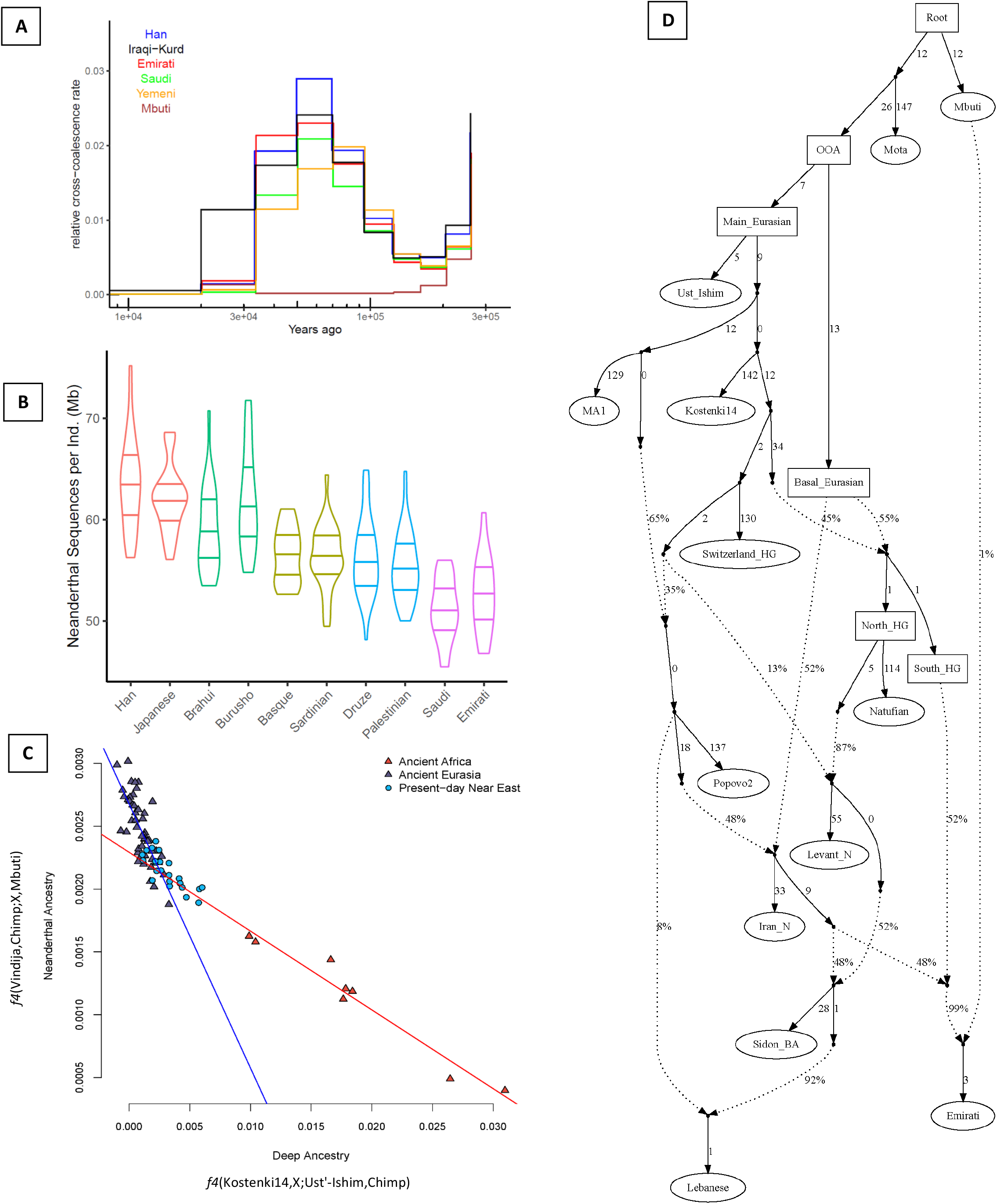
Archaic introgression and deep structure in the Middle East. **A**) Relative cross coalescent rate against Vindija Neanderthal. Note the y-axis range. **B)** Distribution of total length of Neanderthal sequences (Mb) per sample in each population. Horizontal lines depict 25%, 50%, and 75% quantiles. Colors reflect regional grouping. **C)**. Neanderthal ancestry is negatively correlated with a deep ancestry in the Middle East. Two clines explain the depletion of Neanderthal Ancestry in Middle Easterners; one formed by basal Eurasian ancestry and the other is African ancestry. We plot regression lines using the ancient Africans (red) and the ancient Eurasians (blue). **D)** A possible model for the population formation in the Middle East. Populations in ellipses are sampled populations while populations in boxes are hypothetical. Worst f-statistics: (Lebanese, Emirati; Lebanese, Emirati) Z score = −2.83. See Figure S11 for alternative graph models. BA: Bronze Age; HG: Hunter-gatherer.

We then used an identity-by-descent-based method, IBDmix, which directly compares a target population to the Neanderthal genome to detect haplotypes of Neanderthal origin (Chen *et al.*, 2020). We ran IBDmix on our samples and the HGDP dataset, recovering segments totalling ~1.27 Gb that are of likely Neanderthal origin. When comparing the amount of Neanderthal haplotypes that are private to our dataset but not present in other non-Middle Eastern Eurasians, we found only ~25 Mb in total, illustrating that the vast majority of Neanderthal haplotypes in the region are shared with other populations. However, we do find relatively large introgressed haplotypes (~500kb) that are very rare globally, but reach high frequencies in Arabia (Figure S12).

We then compared the average number of total Neanderthal bases per population, and found lower values in Arabia in comparison to other Eurasian populations, including Levantines. The Druze and Sardinians, for example, have similar amounts (average ~56.4 Mb per individual) of Neanderthal ancestry (Figure 3B). In contrast, in Arabia, Emirati.core and Saudi.core have an average of 52.7 Mb and 52.1 Mb Neanderthal ancestry respectively, which is ~8% lower than the Druze and Sardinians, and ~20% less than Han Chinese. Since Emirati.core and Saudi.core have less than 3% of African ancestry, the depletion of Neanderthal ancestry in Arabia cannot be explained by the African ancestry alone. Lazaridis *et al.*, (2014) proposed that a basal Eurasian population, with low-to-no Neanderthal ancestry, had contributed different proportions to ancient and modern Eurasians, reaching ~50% in Neolithic Iranians and Natufians. Since Arabians have an excess of Natufian-like ancestry compared to elsewhere in the Middle East, we find they also carry an excess of basal Eurasian ancestry which will reduce their Neanderthal ancestry. In addition, most modern Middle Easterners carry African ancestry from recent admixture which also contributes to their deep ancestry (relative to the time of a main Eurasian ancestry). We find a negative correlation (Pearson’s r = −0.81, *P* = 2.76e-06) between the increase in deep ancestry and the amount of Neanderthal ancestry in the modern Middle Easterners. When testing all ancient populations we find two clines (Figure 3C) explaining the depletion of Neanderthal ancestry: The first is formed by African ancestry while the second is formed by a Basal Eurasian ancestry in ancient Eurasians. Middle Easterners appear to be affected by both clines since they harbour both ancestries.

### Selection

There is currently a limited understanding of the effects of selection in Arabian populations, with the current hyper-arid climate and a long-term nomad-like subsistence potentially exerting selective pressure for adaptations. To explore this, we searched genome-wide genealogies for lineages carrying mutations that have spread unusually quickly (Speidel *et al.*, 2019) at a conservative genome-wide threshold (*P* < 5×10^−8^). Previous studies identified two correlated variants (rs41380347 and rs55660827), distinct from the known European variant (rs4988235), that are associated with lactase persistence in Arabia (Imtiaz et al. 2007; Enattah et al. 2008). For the Arabian variant rs41380347, we found evidence for strong selection (s = 0.011, logLR = 13.27), similar to, but slightly weaker than, the reported strength of selection at rs4988235 in Europeans (s = 0.016-0.018; Mathieson and Mathieson 2018; Stern et al. 2019). The variant is present at highest frequency in the core Arabian populations: ~50% in Saudis and Emiratis, and at a much lower frequency in the Levant and Iraq (4%). Remarkably, the variant is not present in any Eurasian or African population in the 1000 Genome Project (1KG). We also did not find the variant in published ancient Eurasian whole genomes, including ancient Levantines and Iranians, consistent with a recent origin of the haplotype within the Middle East and subsequent increase in frequency due to selection. We find the variant had a rapid increase in frequency between 9 kya and the present day (Figure 4B). Notably, this period overlaps with the transition from a hunter-gatherer to a herder-gatherer lifestyle in Arabia (Petraglia *et al*. 2020).

**Figure 4.**
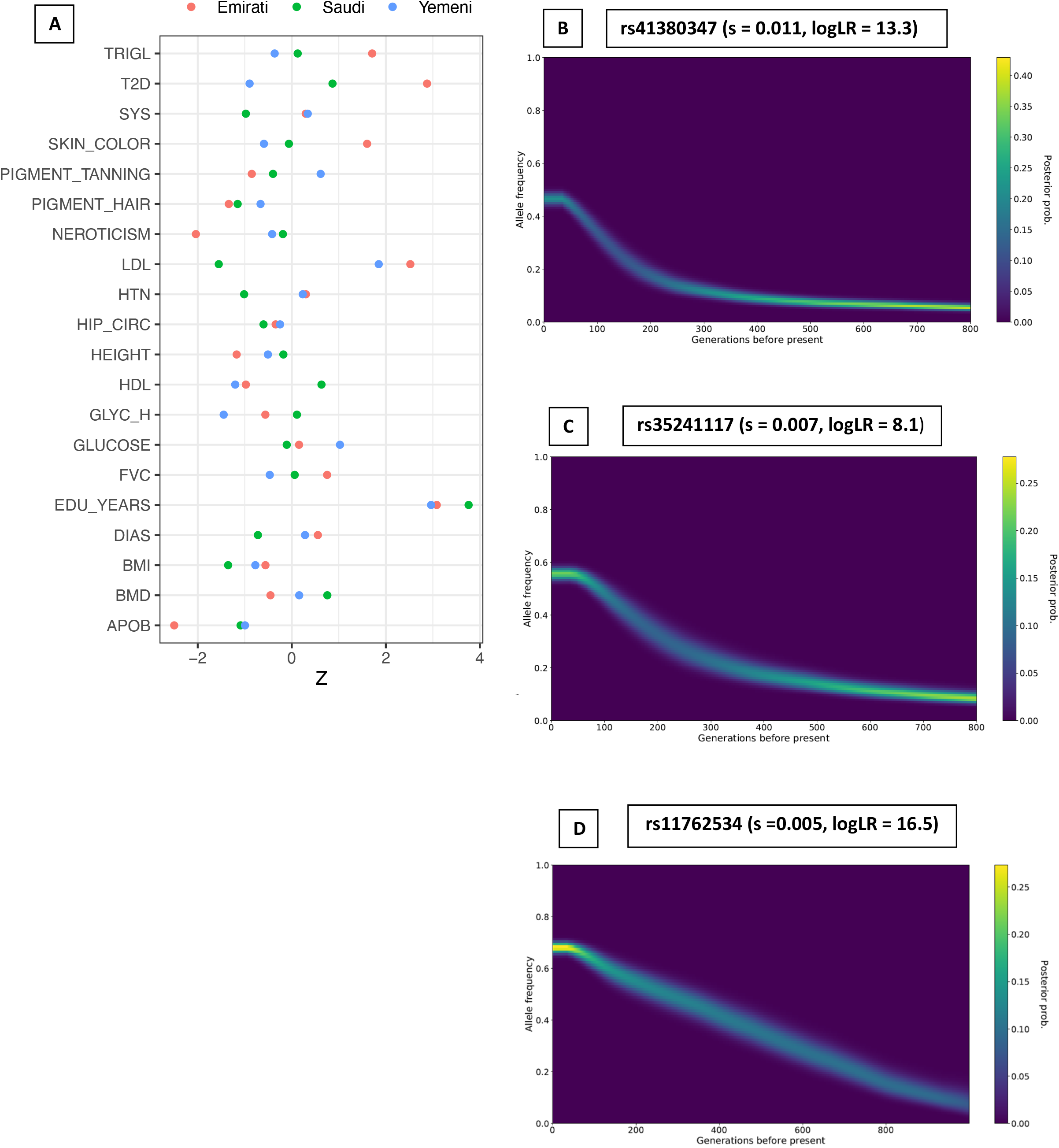
Selection in Arabia. **A)** Testing for recent polygenic selection, over the past 2000 years, on 20 traits within Arabian populations. Asterisks indicate the test is significant after correcting for multiple testing (FDR = 5%). TRIGL: Triglycerides; T2D: Type2 Diabetes; SYS: Systemic Blood Pressure; LDL: Lowdensity lipoproteins; HTN: Hypertension; HIP_CIRC: Hip circumference; HDL: High-density lipoproteins; GLYC_H: Glycosylated haemoglobin; FVC: Forced Vital Capacity; EDU_YEARS: Years of Education; DIAS: Diastolic blood pressure; BMI: Body Mass Index; BMD: Bone Mass Density; APOB: Apoliprotein B **B)** Historical allele trajectory of rs41380347 which is associated with lactase persistence and almost private to the Middle East. s = selection coefficient. **C)** Frequency trajectory of rs35241117, located near *TNKS,* which is present at the highest frequency in Arabia globally and is associated with multiple traits including glomerular filtration rate, bone mineral density, BMI, standing height and hypertension. **D)** Frequency trajectory of rs11762534 which is associated with lymphocyte and neutrophil percentages and prostate neoplasm malignancy and is also present at the highest frequency in Arabia. s = selection coefficient.

We also identified additional variants that show an increase in frequency recently (Figure 4C-D. A variant within *LMTK2*, rs11762534, which is also an eQTL for many genes, displays evidence of selection (s=0.005; logLR = 16.49) and is associated with blood cell percentages and malignant neoplasm of prostate. *LMTK2* encodes a serine/threonine kinase that is implicated in diverse cellular processes including apoptosis, growth factor signalling and appears essential for spermatogenesis in mice (Kawa *et al.,* 2006; Cruz *et al.,* 2019). Outside the Middle East, the variant is highly stratified and is present at the highest frequency in Europeans (1KG, 45%), but we find it at 66% frequency in the Arabian populations. Intriguingly, the variant also shows differentiation in BedouinB (81%), while appearing less frequent in Druze and Palestinians (both ~55%). We additionally looked for strongly differentiated variants between Arabia and the Levant/Iraq (Figure S13). The variant showing the most extreme population branch statistic in Yemenis is rs2814778, where the derived allele results in the Duffy-null phenotype and is almost exclusively found in African populations in the 1000 Genome Project. However, the variant is very common in Yemenis (74%), and decreases in frequency moving northwards in the peninsula (59% in Saudis while reaching 6% in Iraqi-Arabs). We find that across the genome this locus shows the highest enrichment of African ancestry in the Middle East (Methods). As the average amount of African ancestry in Yemenis and Saudis is ~9% and ~3% respectively, the high frequency of this variant appears consistent with positive selection after African admixture. It has been thought that the derived allele protects against *Plasmodium vivax* infection (Miller et al., 1976), which has been historically present in Arabia.

An advantage of using genome-wide genealogies is its power to detect relatively weak selection. We subsequently searched for evidence of polygenic adaptation in Arabian populations across 20 polygenic traits specifically over the past 2,000 years (Methods). For most traits, we find no, or inconclusive, evidence for recent directional selection, including height, skin colour, and BMI (Figure 4A). However a few traits do show evidence, with selection for higher years of education (EduYears) showing the strongest signal consistent across all Arabian populations (*P* = 0.0002 in Saudis). This has also been reported in the British population (Stern *et al.*, 2020); however, the signal was shown to become attenuated after conditioning on other traits, suggesting indirect selection via a correlated trait. In contrast to findings in the British population (Stern *et al.*, 2020), we do not find selection acting on traits such as sunburn, hair color and tanning ability. Within Arabia, the direction of selection on most traits appears to be similar across populations, likely as a result of shared ancestry; however, we note that the current varied environments across the region can potentially cause different recent selective pressures. In Emiratis, we find evidence of selection on variants increasing type 2 diabetes (T2D, *P* = 0.004). This result is intriguing, as the prevalence of T2D in Emiratis is among the highest globally and is partly thought to result from strong recent shift to a sedentary lifestyle (Malik *et al*., 2005). We also find nominal evidence of selection acting to increase levels of low-density lipoproteins (LDL; *P* = 0.01) and decrease levels of Apoliprotein B (APOB; *P* = 0.01) in the same population; but they appear suggestive after adjusting for multiple testing (*P*_*adj*_ = 0.06 at 5% FDR).

## Discussion

In this study we have generated a high-coverage open-access resource from the genetically understudied Middle East region. To our knowledge, this is the first study where the whole population investigated is experimentally-phased, allowing the reconstruction of large and accurate haplotypes. We find millions of variants that are not catalogued in previous global sequencing projects, with a significant proportion being common in the Middle East. A majority of these common variants reside outside of short-read accessibility masks, highlighting the limitation of standard short-read sequencing based studies.

The large number of physically-phased haplotypes allowed us to study population history from relatively old periods (>100 kya) to very recent times (1 kya). We find no evidence that an early expansion of humans out of Africa has contributed genetically to present-day populations in the region. This finding adds to the growing consensus that all contemporary non-African modern humans descend from a single expansion out-of-Africa, quickly followed by admixture with Neanderthals, before populating the rest of the world (Mallick *et al.,* 2016; Bergstrom *et al.,* 2020). We find that Middle Eastern populations have very little Neanderthal DNA that is private to the region, with the vast majority shared with other Eurasians. We demonstrate that Arabian populations have lower Neanderthal ancestry than Levantine, European and East Asian populations and attribute this difference to elevated ancestry from a basal Eurasian population, which did not admix with Neanderthals, in addition to recent African admixture.

By modelling contemporary populations using ancient genomes, we identify differences between the Levant and Arabia. The Levant today have higher European/Anatolian-related ancestry and Arabia having higher African and Natufian-like ancestry. The contrast between the regions is also illustrated by their population-size histories which diverge before the Neolithic and suggest that the transition to a sedentary agricultural lifestyle allowed the growth of populations in the Levant, but was not paralleled in Arabia. It has been suggested that population discontinuity occurred between the late Pleistocene and Early Holocene in Arabia, and that the peninsula was repopulated by Neolithic farmers from the Fertile Crescent (Uerpmann *et al.,* 2010). Our results do not support a complete replacement of the Arabian populations by Levantine farmers. In addition our models suggest that Arabians could have derived their ancestry from Natufian-like local hunter-gatherer populations instead of Levantine farmers.

An additional source of ancestry needed to model modern Middle Easterners is related to ancient Iranians. Our admixture tests show that this ancestry first reached the Levant, and subsequently reached Egypt, East Africa and Arabia. The timings of these events interestingly overlap with the origin and spread of the Semitic languages (Kitchen *et al.*, 2009), suggesting a potential population carrying this ancestry may have spread the language. We find climate change associated aridification events to coincide with population bottlenecks, with Arabians decreasing in size 6kya with the onset of the desert climate while Levantines around the 4.2 kiloyear aridification event. This severe drought has been suggested to have caused the collapse of kingdoms and empires in the Middle East and South Asia, potentially reflected genetically in the signal we identify (Weiss, 2017). Future ancient DNA studies from Arabia are needed to refine the formation of the Arabian populations.

The application of ancestral recombination graphs to reconstruct the evolutionary history of variants offers a powerful method to study natural selection. We refine and identity new signals of selection in Arabian populations. The example of the lactase persistence associated variant, which during the past few thousand years increased to a frequency reaching 50% and is almost absent outside the region, demonstrates the importance of studying underrepresented populations to understand human history and adaptations. Our results indicate that polygenic selection might have played a role in increasing the frequency of variants that were potentially beneficial in the past, but today are associated with diseases such as T2D. We find few signals of polygenic selection in Arabian populations, which may be a consequence of their long-term small effective population sizes which will theoretically reduce the strength of selection. We also note that Middle Eastern populations are among the most understudied populations included in GWAS (Sirugo *et al.*, 2019), which limits the analysis of polygenic traits. Our study and the recent establishment of national biobanks in the region are a step forward to reduce these disparities and offer an exciting opportunity to explore, in the future, complex and disease traits in the Middle East.

## Acknowledgements

We are grateful to Matthew Hurles, Richard Durbin and members of the Tyler-Smith and Martin groups for constructive discussions and comments. We would particularly like to thank the participants for donating samples for this study. We acknowledge the Department of Forensic Science and Criminology at Dubai Police GHQ for allowing us to use their facilities. We thank Faisal Al-Hedeithy, Parwar Hamad, Tariq Zeyad and Mohamed Naji for help with sample collections. M.A.A. was supported by the Government of Dubai - Dubai Police GHQ. C.T.-S and Y.X. were supported by Wellcome grant 098051. P.H. was supported by Estonian Research Council Grant PUT1036.

## Author contributions

M.A.A., Y.X. and C.T-S. conceived this study. M.A.A. and M.H. designed and performed the analyses with contributions from P.H. M.A.A., M.H. Y.X. and C.T-S interpreted the results with input from H.C.M. R.A.L coordinated sample collection and extraction. S.A.T assisted in study design. M.A.A. and M.H. wrote the manuscript. Y.X. and C.T-S. supervised the work. All authors approved the final version of the paper. All authors declare no conflict of interest.

## Data availability

Raw read alignments are available from the European Nucleotide Archive (ENA) under study accession number xxxx. Phased VCFs are available on xxxx.

## Notes

### Competing Interest Statement

The authors have declared no competing interest.

### Summary of Updates

Updated analysis.

