## Supplementary Information for "The Genomic History of the Middle East"

### Methods

##### Sample Collection, Extraction and Sequencing

This study was approved by Dubai Scientific Research Ethics Committee (DSREC-SR-02/2018_01) and by the Wellcome Sanger Institute Human Materials and Data Management Committee (HMDMC 18/026). All individuals who donated samples for this project were interviewed and provided consent for participation. We use the term 'Arabian' in this study to refer to samples from the Arabian Peninsula (Emirati, Saudi and Yemen), Levantine for Syrians and Jordanians, and Iraqi-Arabs and Iraqi-Kurds for samples from Iraq. Saliva samples were collected using the Oragene DNA kits (OG-600) and DNA was subsequently extracted using the Qiagen MagAttract HMW kit (Cat No. 67563). Fragment sizes and quality were determined through a pulsed-field capillary electrophoresis (Femto Pulse system). Libraries were prepared using 10X Genomics Chromium kits and each sample was sequenced in a separate lane on a HiSeq X instrument.

##### Samples Processing and Quality Control

We ran the Long Ranger pipeline (version 2.2.2, using GATK v3.7) to process raw fastq files into phased BAM files and phased VCF files. The average sequencing depth for all samples was 32x, median 31x, calculated using covstats (https://github.com/brentp/goleft/) on the phased BAMs. We assessed the quality of each sample using indexcov (Pedersen et al., 2017) which exploits read depth information from BAMs to identify potential chromosomal abnormalities and quality issues. For each VCF we assessed the number of phased variants through summary.csv file output by Long Ranger, we found on average ~98% of variants to be physically-phased in each sample. For each VCF we assessed the quality of variant calls output by Long Ranger using the QUALITY filter for each variant. The pipeline uses the haplotype structure informed by the physical phasing to tag variants that are likely to be false positives, as each haplotype can only have one allele. We find that variants that do not have a PASS quality label tend to have low Ts/Tv values, suggesting they contain false positives. We used a stringent filter by setting all non PASS variants to missing and merged all samples using bcftools v1.9 merge option -0. We removed variants that show excessive heterozygosity as calculated by bcftools ExcHet tag (< 1e-6) and then set any variant with genotype quality (GQ) < 20 and variants located in regions more than twice the average sample depth to missing. The final dataset composed of 23.1 million single nucleotide variants (SNVs). The Ts/Tv after quality control was 1.97, and remained consistent throughout different allele frequency bins suggesting that the variants are of high quality. We then examined possible relatedness in our dataset using the --genome option in plink-v1.9 (Chang *et al.,* 2015) calculated using a linkage disequilibrium pruned set of 834k biallelic snps (minor allele frequency >5%, missingness <2%, --indep-pairwise 50 5 0.5). We excluded one sample from a pair showing PI_HAT> 0.15, leaving 136 samples for analysis.

##### Chromopainter/fineSTRUCTURE, SOURCEFIND and GLOBETROTTER.

We combined our dataset with published modern global populations extracted from the available curated dataset from the Reich Lab (v37.2) https://reich.hms.harvard.edu/downloadable-genotypes-present-day-and-ancient-dna-data-compiled-published-papers, (samples from the curated dataset analysed in this study were published in Patterson *et al.*, 2012 and Lazaridis *et al.*, 2016), in addition to Ethiopian populations from (Pagani *et al.*, 2015). Variants were lifted over to GRCh38 using picard (v2.18.26, https://broadinstitute.github.io/picard/). We used a minor allele frequency filter >4%, excluded variants with >5% missingness and variants that show departure from Hardy-Weinberg equilibrium (< 1e-10). To avoid bias from different phasing methods, as advised by the authors of the Chromopainter/FineSTRUCTURE software, we discarded for this specific test the physical phasing from our samples and phased the merged dataset with Eagle v2.4.1 (Loh *et al.,* 2016) using the 1000 Genomes Project phase 3 panel (1000 Genomes Project Consortium *et al.*, 2015). We then ran the

Chromopainter/FineSTRUCTURE pipeline v4.1.1 using ~400K variants (Lawson *et al.,* 2012). We initially ran the pipeline on a total of 517 samples to identify homogeneous populations and excluded outliers for the subsequent SOURCEFIND (Chacón-Duque *et al.*, 2018) and GLOBETROTTER (Hellenthal *et al.*, 2014) analysis, as recommended by the authors. We also ran the pipeline on a limited set of 303 Middle Eastern samples to look at the region in more detail (Figures 1D and S2A). Finally, we also ran the pipeline only on our dataset, once using 474K high-quality polymorphic variants we identified previously in the HGDP (within the strict mask and have GATK high variant quality score recalibration) and second using a larger number of variants (2.2 million) identified within this study (Figure S2B). Using 400K variants (Figures 1D and S2A) fineSTRUCTURE could not distinguish Saudi and Yemeni samples, however by using a larger number of variants they became more distinguishable (Figure S2B). We ran the pipeline twice for each dataset to assess variability and found generally consistent results. Based on the output, we divided the self-labelled populations into representative 'core' subpopulations that show limited-to-no recent admixture.

To investigate potential sources of admixture, we used SOURCEFINDv2 (Chacón-Duque *et al.*, 2018), a haplotype-based method that represents a population as a mixture of surrogates. We chose donor populations defined by the previous fineSTRUCTURE results and in some cases combined some populations that clustered together. We define 'core' and 'mix' samples, 'mix' are a set of samples that cluster together that are not 'core', i.e. potentially admixed. We ran ChromopainterV2 based on a set of the following donor and recipient populations ("_" illustrates that populations were combined):

<u>Donor</u>: Assyrian_Armenian_Georgian, Bantu_Kenya_Luhya_Luo, BantuSA, Bulgaria_Albania, Chechen_Kum_Lez, Esan_Yoruba, Gumuz, Iranian, Iranian.Bandari, Kalash, Khomani_San, Kyrgyz, Lebanese, Makrani_Brahui_Balochi, Malta_Sicily, Mbuti_Biaka, Mende_Mandenka, Pathan_Sindhi, Punjabi_Burusho, Sardinian, Saudi.core, Uzbek_Turkmen.

<u>Recipient</u>: Donors and Saudi.Mix, Emirati.core, Emirati.Mix1, Yemeni, Jordanian, Iraqi Arab, Iraqi Kurd, Syrian and Omani

We also repeated the previous step replacing Saudi.core from the donor and including Emirati.core instead. We ran SOURCEFINDv2 using default parameters, 100 separate times setting the previous donor populations as 'surrogates' and recipient populations as 'targets'. For each result we took the sample with the highest posterior probability and averaged the 100 samples.

Notably, in Eastern Arabia, we find that the Emiratis formed multiple distinct groups. SOURCEFIND infers one of these Emirati clusters to be similar to Saudis, another is closest to coastal Iranians. This substructure in the Emirati population is visible in Figure S2B, where a subset of Emirati samples cluster closer to Levantines rather than other Arabians. We excluded the coastal Iranian-like Emirati cluster from Figure 1D and S2A, but they are shown in Figures S2B.

We subsequently used fastGLOBETROTTER, a newer implementation of GLOBETROTTER which was originally described in Hellenthal *et al.,* 2014, using the parameters (prop.ind: 1, bootstrap.date.ind: 1, null.ind: 1, with all remaining parameters default) including only as surrogates populations that contributed >1% ancestry in the previous SOURCEFIND step (Figure S5). The idea of including Saudi.core (or Emirati.core) as a donor was to have a representative source from Arabia. As having a target population as a donor and recipient can mask signals due to self-copying, we re-ran the step substituting Saudi.core with Emirati.core to investigate the Saudi population. Moreover, as Emirati.core and Saudi.core are themselves similar, we also ran fastGLOBETROTTER excluding both Emirati.core and Saudi.core as surrogates, instead including Lebanese. We also tested for admixture using MALDER (Loh *et al.*, 2013; Pickrell *et al.*, 2014). We included 590k variants using 6 references:

(Luhya in Webuye, Kenya (LWK.SG);Yoruba;Druze;Iranian;Indian Telugu in the UK (ITU.SG); Punjabi in Lahore, Pakistan (PJL.SG) and ran using default parameters (Table S2).

**Demographic History**

We leveraged the physical-phasing in our dataset using RELATE v1.1 (Speidel *et al.*, 2019) to examine effective population size and separation history. We generated genome-wide genealogies using 54 physically-phased haplotypes based on "core" samples curated from fineSTRUCTURE results. We limited analysis to regions within the genome accessibility mask described in Bergstrom *et al.,* 2020 and set unphased variants to missing (i.e. excluded from analysis). We then converted phased VCFs to the haps/sample file format using the RelateFileFormats script (part of the Relate package) and prepared the input files using PrepareInputFiles.sh. We supplied the human ancestor sequences downloaded from ftp://ftp.ensembl.org/pub/release-100/fasta/ancestral_alleles/ to polarize variants as ancestral or derived. We then ran Relate with with options -m 1.25e-8 -N 30000 using the HapMap genetic map supplied with Eaglev2.4.1 (genetic_map_hg38_withX.txt.gz) then used the output in the EstimatePopulationSize.sh script with options -m 1.25e-8 --years_per_gen 29.

We were concerned that the decrease in population size we find around 4-5kya could be a result on recent consanguinity, which is common in the Middle East. Our runs of homozygosity (ROH) analysis show that some Arabian samples in particular have large blocks of homozygosity which is consistent with recent consanguinity (Figure S15). To test this, we repeated the population size analysis, firstly including samples with high total sum of ROH (sROH > 50 Mb; minimum ROH size of 1Mb), secondly including samples with relatively low sROH (< 50 Mb) and thirdly by choosing one haplotype per individual, instead of two haplotypes. This will remove the effect of recent consanguinity, as we also removed any related samples as discussed in the 'Samples Processing and Quality Control' section. We show the results of these tests in Figure S16A. Including samples with high total of ROH amplifies the population reduction in the past 4ky and then a modest recovery in the last 1ky is observed. Samples with relatively lower ROH show a similar history, but the recent decrease in size is more attenuated and a stronger recovery is observed in the last 1ky. The single haplotype curves show that the recent recovery is slightly older and begins at 2kya and results in a larger increase in population size in comparison to the previous two curves. The second bottleneck appears in all curves. We also repeated the analysis on single haplotypes separately for both Levantines/Iraqis and Arabians and find similar results. Figure 2A in the main text refers to the single haplotype analysis. For the separation history analysis Figure 2C we used the low sROH diploid samples shown in Figure S15.

As we find a divergence in population size between Arabia and the Levant before 10kya, i.e. before the Neolithic era, which has potential relevance regarding the theory of the peopling of Arabia by Levantine Neolithic farmers, we reran the analysis using another method (MSMC2 v2.1.1; first described in Schiffles and Durbin, 2014 with later version MSMC2 published in Wang *et al.*, 2020) to check for concordance with the results from Relate. We used 4 haplotypes from 4 individuals (1 haplotype per individual) per population. The results from MSMC2 agree with Relate, the divergence in size starts before 10kya (Figure S16A).

For the separation history analysis with global populations, we downloaded the Mbuti, Sardinian, and Han Chinese physically-phased samples from the HGDP (2 samples per population, Bergstrom *et al.*; 2020). Since the published data used an older version of Long Ranger, we recalled the samples using v.2.2.2 to be consistent with our dataset. We filtered the VCFs as described above for our dataset. We used MSMC2 v2.1.1 to infer split times between our populations and the HGDP samples using 8 haplotypes for each comparison (4 haplotypes from each population). MSMC2 was run using the --skipAmbiguous option, to calculate coalescent rates within and between populations, restricted to the genome accessibility mask described in Bergstrom *et al.,* 2020. We used a generation time of 29 years and a mutation rate of 1.25e-8 to scale the results. We then used MSMC-IM (Wang *et al.,* 2020) on the output of the previous step to infer migration rates from coalescent rates using default parameters (Figure S10). We excluded from analysis migration rates when the cumulative migration probability reached over 0.999, as suggested by the authors.

**Archaic Admixture**

We used IBDMix (Chen *et al.,* 2020) to call Neanderthal segments in a merged dataset of our samples with the HGDP. We downloaded the high coverage Altai Neanderthal (Prüfer *et al.*, 2014) and Denisova (Meyer *et al.*, 2012) VCFs from http://cdna.eva.mpg.de/neandertal/Vindija/VCF/. We excluded modern populations with less than 10 samples as suggested in Chen *et al.,* 2020 and followed the filtering steps they previously described: We removed sites that lie within segmental duplications downloaded from (http://hgdownload.cse.ucsc.edu/goldenPath/hg38/database/genomicSuperDups.txt.gz), removed variants that are CpG, restricted analysis to the previously described HGDP accessibility mask, in addition to the archaic genome accessibility masks downloaded from https://bioinf.eva.mpg.de/altai_minimal_filters/. We removed singletons, variants that deviate from Hardy-Weinberg equilibrium (< 1e-10), show excessive heterozygosity (< 1e-8), and only included biallelic snps. As performed in Chen *et al.,* 2020, we also ran IBDMix to identify 'Denisovan' segments in African populations and masked these regions in non-Africans as they are likely to be enriched for incomplete lineage sorting. We filtered the remaining segments using a minimum size threshold of 50kb and LOD score higher than 4.

We also ran Sprime (Browning *et al.,* 2018) on a similar dataset as above but without excluding CpG sites, regions of segmental duplications and the archaic genome accessibility masks. As Sprime requires non-missing genotypes, we removed variants that were >5% missing and imputed the remaining missing variants using Eaglev2.4.1 (Loh *et al.,* 2016). We set all non-Middle Eastern samples from the HGDP as outgroup (768 samples) and ran Sprime for each Arabian population. We filtered the output using a score threshold of 150,000.

We applied MSMC2 using 4 haplotypes (2 diploid samples) to examine the separation history between our populations and the high coverage Vindija Neanderthal (Prufer *et al.* 2017) as we did in our previous study (Bergstrom *et al.,* 2020). Briefly, this analysis exploits the fact that the Vindija Neanderthal shows extremely low heterozygosity, which renders much of the genome homozygous and essentially phased. We used the --skipAmbiguous option to exclude sites with unknown phase and ran MSMC2 using the same parameters previously stated.

**Selection**

We used the Relate Selection Test in RELATE v1.1 (Speidel *et al.,* 2019) to look for lineages that spread faster than competing lineages. At every site, all variants are required to be phased to perform this test and our relatively stringent quality control may set some variants as missing (for e.g. if a sample had a duplication at a locus, variants within the region will be set as missing because of our depth filter), so we relaxed the depth filter for this test. In addition, as the 10X linked-read technology phases ~98% of variants, we used BEAGLEv4.0 (Browning and Browning, 2014) to statistically phase the remaining variants using the gtgl option only for this specific selection test. We set the option usephase=true to take into account the physical-phasing already provided in the VCF. We first ran RELATE v1.1 on 272 haplotypes using the same parameters described in the demographic history section. We extracted the genealogies of the Arab.core samples and used the output of the previous step as input to the DetectSelection.sh script accounting for the population history of the populations using the option -m 1.25e-8 --years_per_gen 29. From the resulting .sele file, we extracted p-values from the column "when_mutation_has_freq2" which tests for evidence of selection over the lifetime of a particular variant. We were conservative and only included variants that show significance at a "genome-wide threshold" of $P < 5e\text{-}8$. This test for significance has been shown to be well calibrated (Spiedel et al., 2019), but to even further refine and understand the evolutionary history of the variant we used CLUES (Stern *et al.*, 2019; https://github.com/35ajstern/clues). From the output of Relate above, we ran the SampleBranchLengths.sh script to sample branch lengths from the posterior in order to account for uncertainty. We ran 100 samples (--num_samples 100) using a mutation rate of 1.25e-8 and accounted for the population size history by supplied the .coal files from the previous step. We then ran CLUES (inference.py script) with the option --coal {.coal file} to again account for population size changes. We fine-mapped variants using the likelihood ratio statistic produced by CLUES as suggested by Stern *et al.,* 2019 and focused on variants that show moderate to strong selection (s > 0.005). We used the plot_traj.py script to plot the results.

We find a signal of strong selection at rs35241117 (Figure 6C, s = 0.007, logLR = 8.1). This variant shows the highest global frequency in Saudis and Yemenis (~60%), and is associated with a number of metabolic, skeletal and immunological traits, including glomerular filtration rate, diuretics, hypertension and BMI. rs35241117 lies outside a ~400 kb haplotype that has recently been suggested to be under selection in Kuwaitis and Saudis (Eaaswarkhanth *et al.*, 2020), but is in moderate LD ($r^2 = 0.51$) with it. It seems likely that the selection signal observed by Eaaswarkhanth *et al.*, whose dataset is composed of array data, is actually linked to rs35241117, which based on our analysis underestimates the strength of selection by almost a half

To test for polygenic selection, we used PALM (Stern *et al.*, 2020; https://github.com/35ajstern/palm). We avoided the use of GWAS summary statistics that were calculated from meta-analysis, due to the potential effect of uncorrected population stratification. We extracted GWAS summary statistics performed on the UK BioBank (UKBB; (Bycroft *et al.*, 2018) downloaded from (https://data.broadinstitute.org/alkesgroup/UKBB/UKBB_409K/ - (Gazal *et al.*, 2018); https://data.broadinstitute.org/alkesgroup/UKBB/LTFH/sumstats/ - (Hujoel *et al.*, 2020) ; and from the Neale lab Imputed v3 dataset http://www.nealelab.is/uk-biobank/ ). These statistics were nominally corrected for population structure using either a family history-based approach, fixed PCs, or a linear mixed model. For each trait investigated, we split the genome into 1,700 approximately independent blocks (Berisa and Pickrell, 2016) and selected the variant with the lowest p-value within each block for analysis. We chose to be conservative by only including blocks with variants that were genome-wide significant ($P < 5e\text{-}8$), as potentially uncorrected population structure is not expected to produce such highly significant values. Moreover, the method we use, PALM, has been shown to perform well even with some uncorrected GWAS stratification (Stern *et al.,* 2020). Variants were filtered for minor allele frequency > 5%, Rsq > 0.5, INFO score > 0.8, and excluded indels. For each variant passing the previous thresholds, we sampled branch lengths as done for CLUES and estimated selection likelihoods with the lik.py script using the options --coal {.coal file} to account for population size changes and options --K 1 --kappa 3 to specifically test for selection over the past 2000 years, assuming a generation time of 29 years. We then ran palm.py to test for polygenic selection using the option --B 1000 (number of bootstraps). To further explore the choice of significance threshold, and potential effects of uncorrected population structure (variants passing a higher significance threshold are likely to be less biased by uncorrected population structure), on the results, we repeated the analysis using more stringent significance thresholds ($P < 1e\text{-}8$ and $P < 5e\text{-}9$; which will drop blocks not passing the threshold), and found similar values (Figure S14).

**PHEWAS and eQTL analysis**

We used the Phewas search option in the GWAS atlas (Watanabe *et al.*, 2019) and Gene Atlas (Canela-Xandri et al., 2018) to look for trait associations with variants that show evidence of selection. We used the GTEx portal (GTEx Analysis Release V8; The GTEx Consortium, 2020) to look for eQTL associations.

**Runs of homozygosity analysis.**

We used plinkv1.9 (Chang *et al.,* 2015) to first filter and prune our dataset using the options: --geno 0.05 --indep-pairwise 50 5 0.5 --maf 0.05 and subsequently identified runs of homozygosity (ROHs) using the option --homozyg with all other options kept as default. We restricted the analysis to the strict mask defined previously. This identifies ROHs ≥ 1 Megabase (Figure S15).

**Y-chromosome and Mitochondria analysis**

Y-chromosomal data of 79 males from the current study were complemented by 46 samples from Haber et al. 2019 and 1208 samples from Hallast et al. 2020. Genotype calling, filtering and Y haplogroup prediction are described in detail in Hallast et al. 2020. Additionally, 11 samples with >4% of missing data from Haber et al. 2019 were removed from the final analysis. After filtering a total of 1322 samples and 10,194,410 sites remained, including 90,810 variant sites.

All 79 males from the current study, 35 males from Haber et al and 332 selected informative males in the context of the study from Hallast et al. 2020 were used to estimate the ages of internal nodes in the Y phylogeny using the coalescence-based method implemented in BEAST (v1.8.4, Drummond and Rambaut 2007). This dataset of 446 samples contained 10,194,410 sites, including 49,728 variant sites. A starting maximum likelihood phylogenetic tree for BEAST was constructed with RAxML (v8.2.10, Stamatakis 2014) with the GTRGAMMA substitution model using variant sites. Markov chain Monte Carlo samples were based on 277 million iterations, logging every 1,000 iterations and the first 10% of iterations discarded as burn-in. LogCombiner was used to combine 20 independent runs. The HKY substitution model accounting for site heterogeneity (gamma), a constant-sized coalescent tree prior and strict clock with a substitution rate of 0.76 $x10^{-9}$(95% confidence interval: $0.67x10^{-9}$ to $0.86x10^{-9}$) single nucleotide mutations per bp per year (Fu et al. 2014) was used. A prior with a normal distribution based on the 95% confidence interval of the substitution rate was applied. Only the variant sites were used, but the number of invariant sites was defined in the BEAST xml file. A summary tree was produced using TreeAnnotator (v1.8.1) and visualised with the FigTree software (Figure S9; http://tree.bio.ed.ac.uk/software/figtree/).

We created a per sample all-site VCF of the mitochondrial reference sequence contig from each sample BAM file using the command "bcftools mpileup -C 50 -Q 20" and called haplogroups using haplogrep-v2.1.25 (Weissensteiner et al., 2016). We then aligned the sequences using MUSCLE-v3.8.31 (Edgar, 2004), cleaned the alignments using Gblocks-v0.91b (Castresana, 2000), and ran ModelTest-NG (Darriba et al., 2020) to select the best fitting nucleotide substitution model (TIM1+I+G4). We used RAxML-NG-v.0.9.0 with options "--support --bs-trees --tree" to construct a maximum-likelihood phylogenetic tree (Kozlov et al., 2019). We visualized the tree using FigTree (Figure S17).

**Local Ancestry Deconvolution**

We used RFMix v2.03 (Maples et al., 2013) to identify African haplotypes within our dataset. We used 105 samples from the HGDP as references: 41 Druze and 64 Africans. The samples were chosen based on a previous ADMIXTURE run (Bergstrom et al., 2020) and outliers were excluded (i.e. Druze that show relatively high African ancestry, or Africans that show relatively high Eurasian ancestry). RFMix was run using the option -e 5 for 5 EM iteration steps with all other options set as default.

**Ancient DNA dataset**

We merged our new data with published ancient data extracted from a previously curated dataset available from the Reich lab (v42.4) (Allentoft et al. 2015; Antonio et al. 2019; de Barros Damgaard et al. 2018; Feldman et al. 2019; Fregel et al. 2018; Fu et al. 2014; Gamba et al. 2014; Harney et al. 2018; Jones et al. 2015; Lazaridis et al. 2017; Lazaridis et al. 2016; Lipson et al. 2017; Gallego Llorente et al. 2015; Mathieson et al. 2015; Mathieson et al. 2018; Narasimhan et al. 2019; Olalde et al. 2018; Olalde et al. 2019; Prendergast et al. 2019; van de Loosdrecht et al. 2018; Schuenemann et al. 2017; Villalba-Mouco et al. 2019; Mittnik et al. 2018; Gunther et al. 2015), we also extracted modern individuals from worldwide populations genotyped on the Human Origins array (Patterson et al. 2012, Lazaridis et al. 2014, Lazaridis *et al.,* 2016). We added ancient and modern Levantines (Haber *et al.*, 2017; Agranat-Tamir et al. 2020), and modern Egyptians and Ethiopians (Pagani *et al.* 2015). We converted the coordinates of the published data to the human genome assembly GRCh38 using the LiftOver tool from the UCSC Genome Browser http://genome.ucsc.edu/ and used bcftools v1.9 with command "bcftools mpileup -T -q30 -Q30 | bcftools call -c" to genotype our samples for positions found in the ancient DNA data. We merged the datasets using the mergeit program available from the EIGENSOFT package v7.2.1 with options docheck: YES and strandcheck: YES filtering sex-linked and triallelilic SNPs and sites that were outside the accessibility mask (Bergstrom *et al.,* 2020). The final dataset included 4,150 individuals and 443,885 SNVs. For the LD-decay tests we excluded the samples genotyped on the Human Origin array which increased the available number of SNVs to 815,350. We add the suffix ".HO" to the names of the published modern Middle Eastern populations to differentiate them from our new samples.

**Principal component analysis and model-based clustering**

We computed a PCA using smartpca v16000 from the EIGENSOFT package (Patterson et al., 2006) with parameters numoutlieriter: 0, lsqproject: YES, autoshrink: YES and using only variation in modern populations selected to represent genetic diversity in Central Asia, the Middle East, Europe, and East Africa (Figure 1B and S3).

We ran ADMIXTUREv1.3 (Alexander et al., 2009) in an unsupervised mode from K=6 to K=20 using ~80,000 transversions in our dataset which we randomly subsetted to ≤ 10 individuals per modern population and ≤ 20 individuals per ancient population. We show K=10 which had the lowest cross-validation error compared with other K values, and also show K=17 when Natufians and Neolithic Anatolians formed separate ancestral components (Figure S6). We ran DyStruct (Figure 1C; Joseph et al., 2019) on the same dataset with default arguments across nine time points binned as follows (in years ago): 14,500-10,000; 10,000-8000; 8000-6000; 6000-5200; 5200-5000; 5000-3000; 3000-1400;1400-200; and present-day.

***f4* statistics, qpAdm, qpGraph and MALDER**

From the ADMIXTOOLS package (Patterson *et al.*, 2012) we used qpDstat v755 with parameter f4mode: YES to test the genetic contrast between North and South of the Middle East (Figure S4) and to assess the amount of Neanderthal and Basal Eurasian ancestry. We used *qpAdm* v810 with option allsnps:YES to estimate ancestry proportions in our samples and used *qpGraph* v6450 to draw phylogenetic models that explain the formation of populations in the Middle East.

We used MALDER v1.0 (Loh *et al.* 2013; Pickrell *et al.* 2014) with parameters mindis: 0.005, binsize: 0.0005 and a generation time of 28 years to estimate admixture time related to ancient Iranians from decay of LD. We tested our populations using weights from ancient Levantines and ancient Iranians: Natufians, Levant_N, Levant_ChL, Iran_N (Ganj Dareh), Iran_Tepe_Hissar_ChL, Iran_Seh_Gabi_ChL, and Iran_Hajji_Firuz_ChL. We similarly tested admixture in East Africans but replaced the ancient Levantines references with Gumuz and Yoruba.

### Supplementary Figures

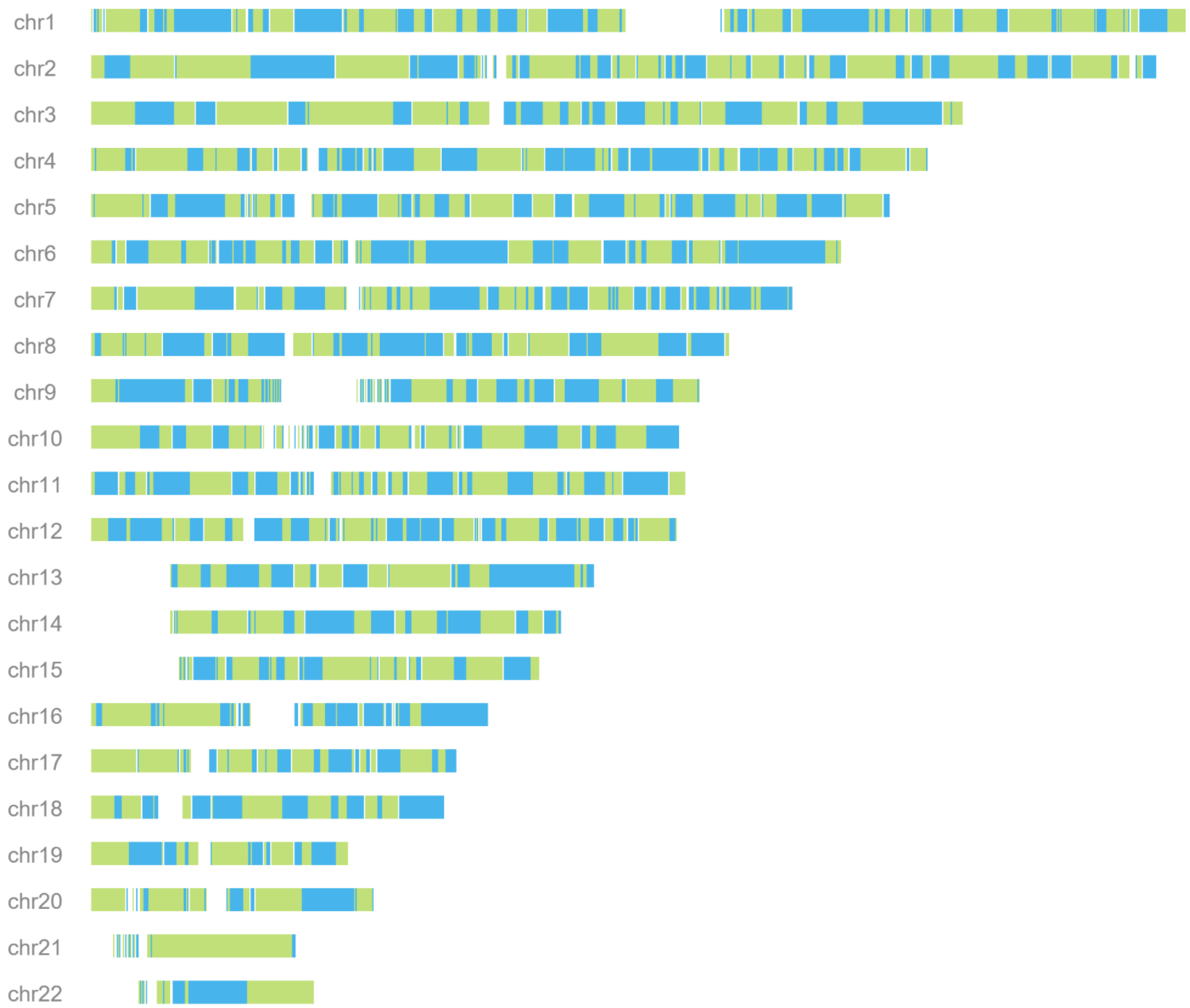


**Figure S1: Physical-phasing using linked-reads.** Example using sample APPG7555924. Physically-phased blocks illustrated using alternating green and blue colors. Longest phase block: 31.9 Mb. N_50 phase block: 5.2Mb.

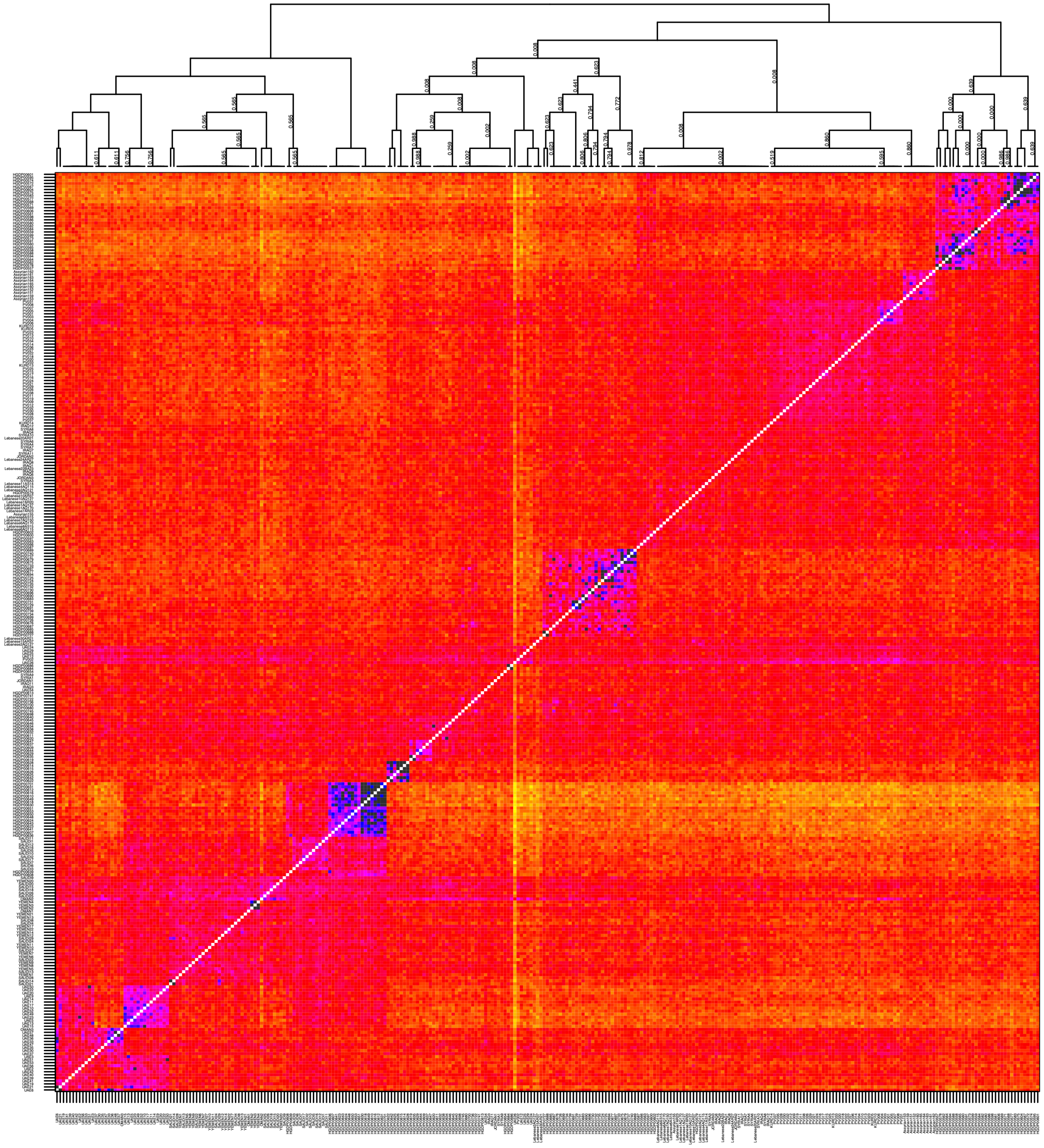

**Figure S2A: Chromopainter co-ancestry matrix of Middle Eastern samples from Figure 1.** Heatmap capped at 200 shared segments (dark segments). Numbers on tree edges represent the posterior assignment probabilities from fineSTRUCTURE, edges with no numbers have a posterior assignment probability = 1. Samples generated from this study are capitalized to distinguish them from other Middle Eastern samples curated from Reich Lab dataset (v37.2).

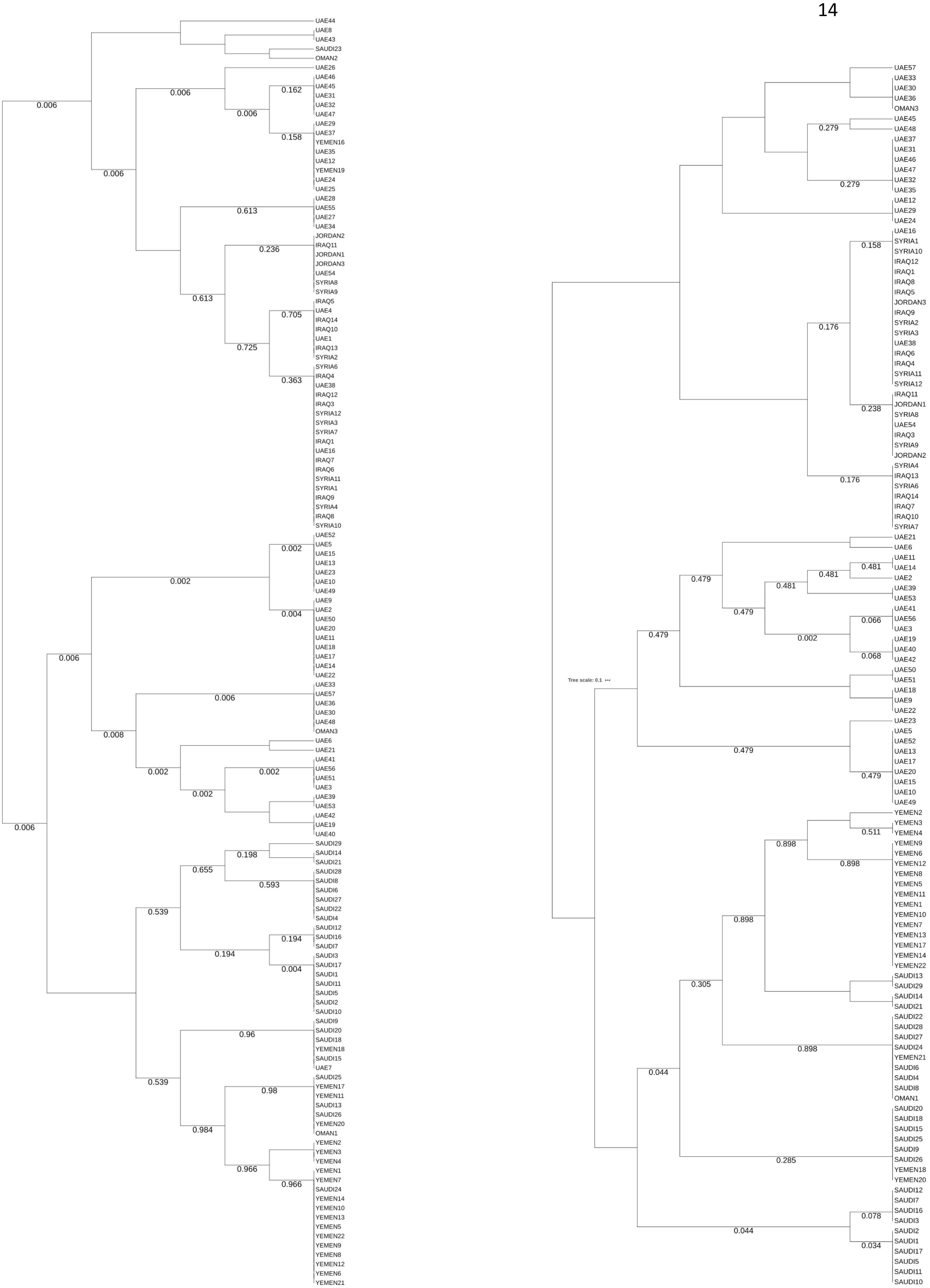

**Figure S2B: fineSTRUCTURE trees of samples generated in this study.** Numbers on tree edges represent the posterior assignment probabilities, edges with no numbers have a posterior assignment probability = 1. **Left:** Using 474K high-quality common (>5%) polymorphic variants identified in the HGDP. **Right:** Using 2.2M variants identified within our dataset excluding 16 potentially recently admixed samples. Note a subset of Emirati samples cluster with Levantine populations; these samples are inferred to have ancestry similar to coastal Iranians (Bandari).

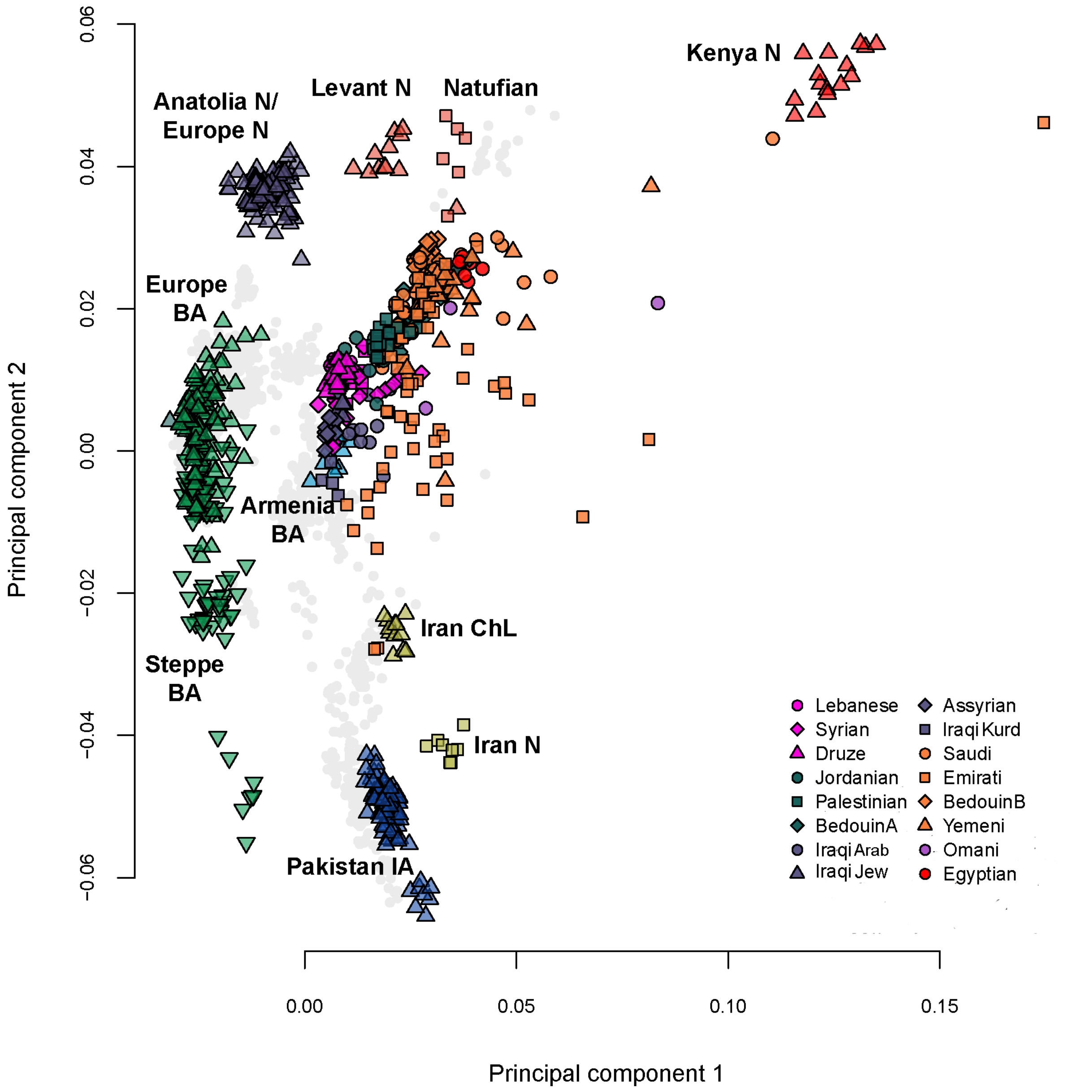


**Figure S3: Principal Components Analysis.** Plot similar to Figure 1 but including recently admixed samples in our dataset (i.e. non-core). Eigenvectors were inferred with present-day populations from the Middle East, North and East Africa, Europe, Central and South Asia (all modern non-Middle Easterners shown as grey points). The ancient samples were then projected onto the plot.

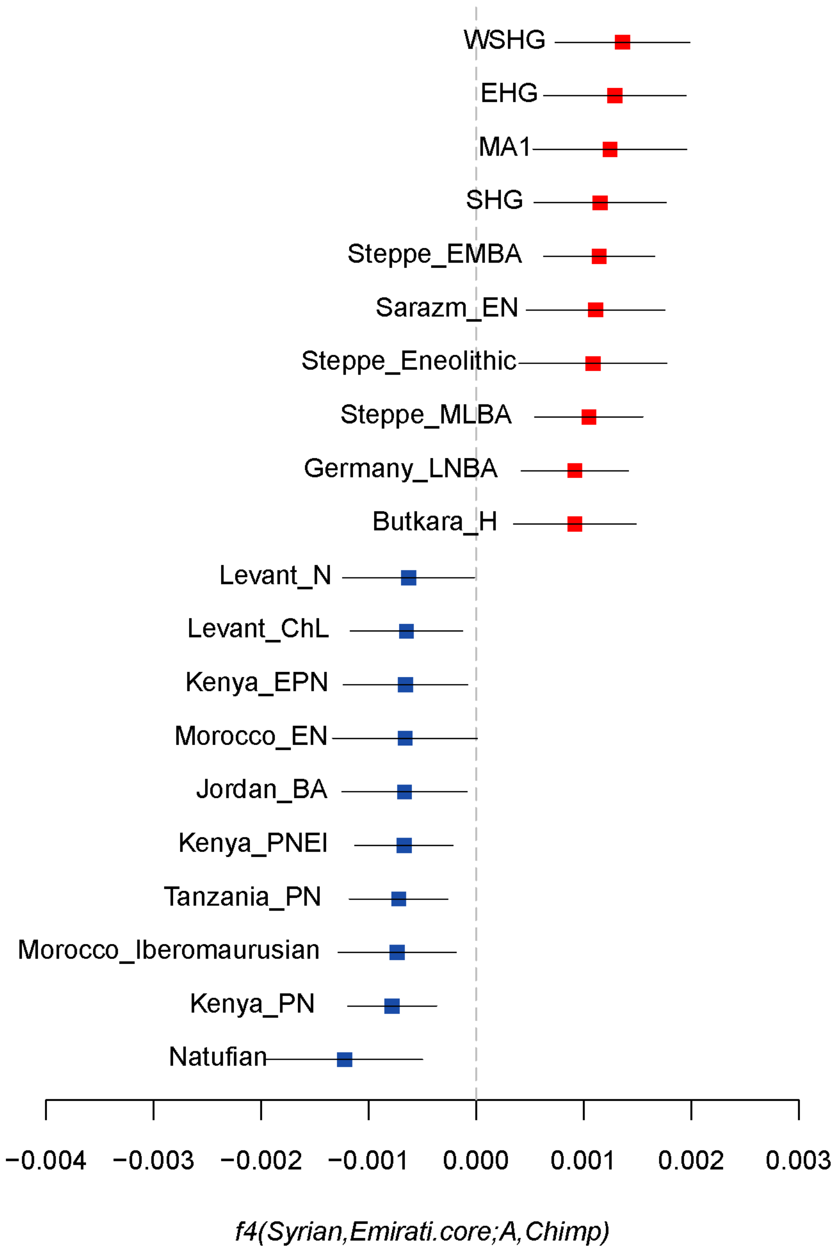


**Figure S4: Genetic contrast between the Levant region and Arabia.** We plot the statistic *f4*(Syrian,Emirati-core;Ancient,Chimpanzee) and ±3 standard errors from results with the 10 lowest (blue) and 10 highest (red) f4-stats.

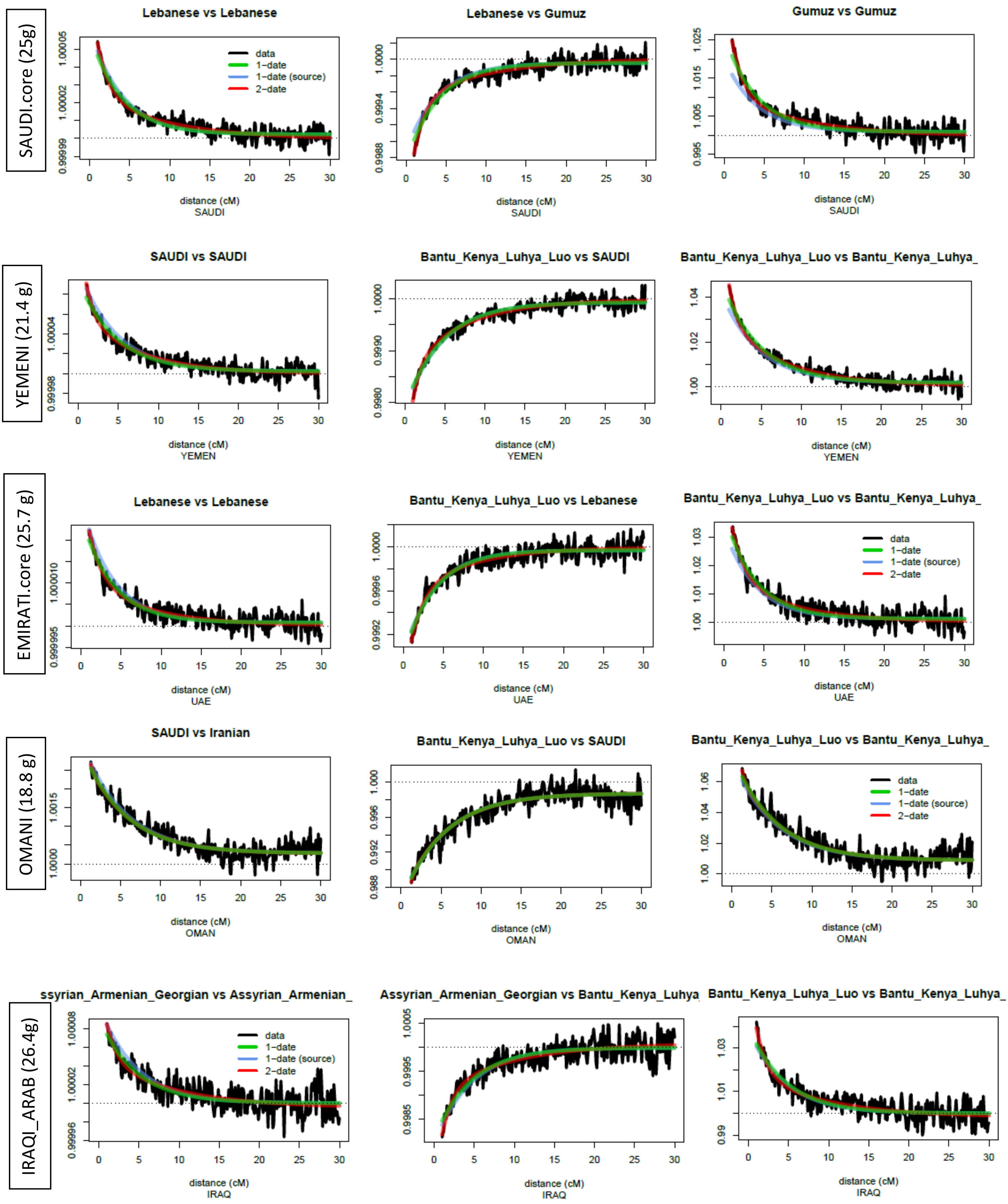

SAUDI.core (25g)
Lebanese vs Lebanese
data
1−date
1−date (source)
2−date
distance (cM)
SAUDI
Lebanese vs Gumuz
distance (cM)
SAUDI
Gumuz vs Gumuz
distance (cM)
SAUDI
YEMENI (21.4 g)
SAUDI vs SAUDI
distance (cM)
YEMEN
Bantu_Kenya_Luhya_Luo vs SAUDI
distance (cM)
YEMEN
Bantu_Kenya_Luhya_Luo vs Bantu_Kenya_Luhya_
distance (cM)
YEMEN
EMIRATI.core (25.7 g)
Lebanese vs Lebanese
distance (cM)
UAE
Bantu_Kenya_Luhya_Luo vs Lebanese
distance (cM)
UAE
Bantu_Kenya_Luhya_Luo vs Bantu_Kenya_Luhya_
data
1−date
1−date (source)
2−date
distance (cM)
UAE
OMANI (18.8 g)
SAUDI vs Iranian
distance (cM)
OMAN
Bantu_Kenya_Luhya_Luo vs SAUDI
distance (cM)
OMAN
Bantu_Kenya_Luhya_Luo vs Bantu_Kenya_Luhya_
data
1−date
1−date (source)
2−date
distance (cM)
OMAN
IRAQI_ARAB (26.4g)
ssyrian_Armenian_Georgian vs Assyrian_Armenian_
data
1−date
1−date (source)
2−date
distance (cM)
IRAQ
Assyrian_Armenian_Georgian vs Bantu_Kenya_Luhya
distance (cM)
IRAQ
Bantu_Kenya_Luhya_Luo vs Bantu_Kenya_Luhya_
distance (cM)
IRAQ

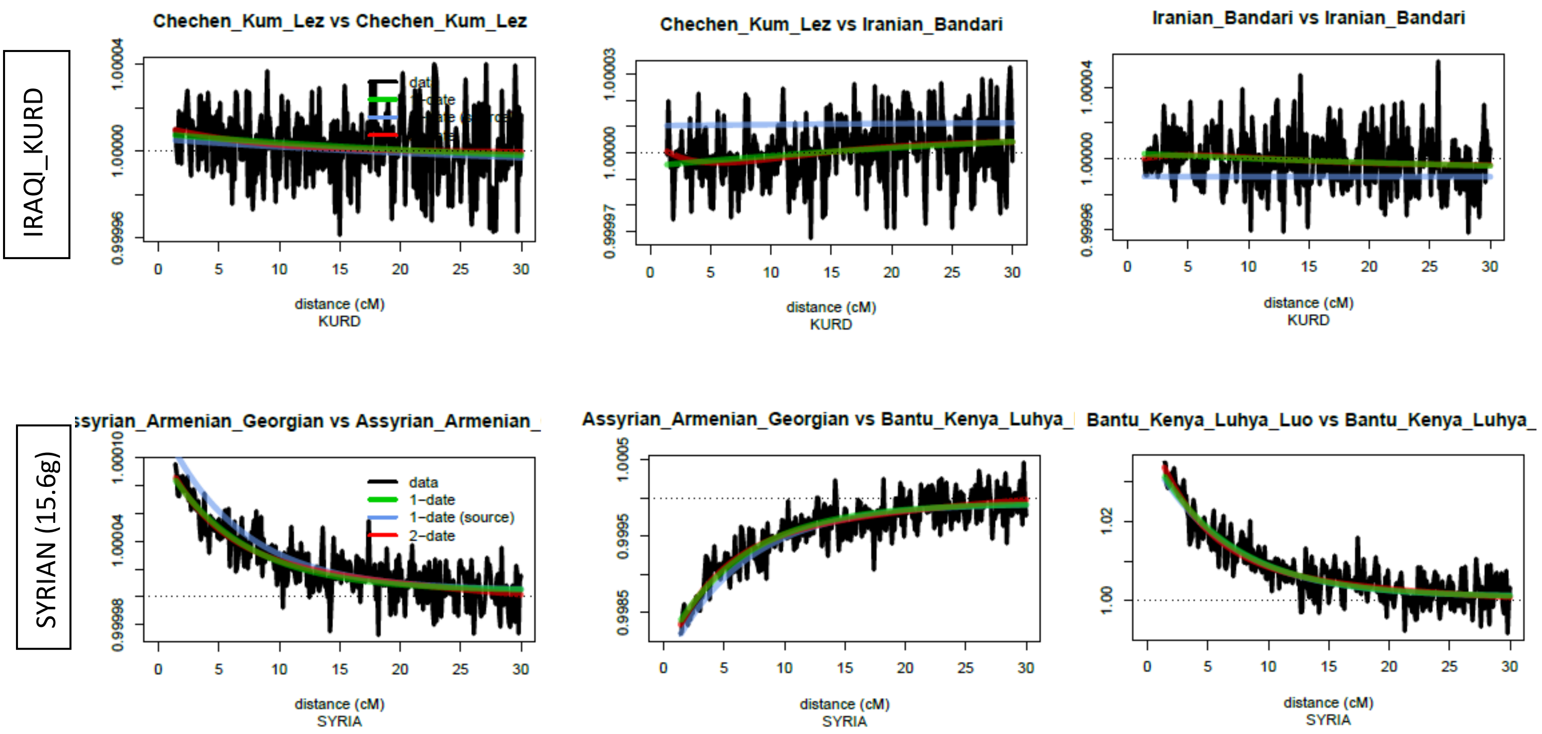


**Figure S5: Admixture using Globetrotter.** Co-ancestry curves showing relative probability of jointly copying two chunks from donors at varying genetic distances. The curves fit an exponential decay (1-date green line, 2-date red line). The positive slope (middle curve) implies that these donors represent admixing sources. The estimated admixture date is illustrated on the left of each figure, g for generations. The Iraqi_Kurds are notable for not showing recent evidence of admixture.

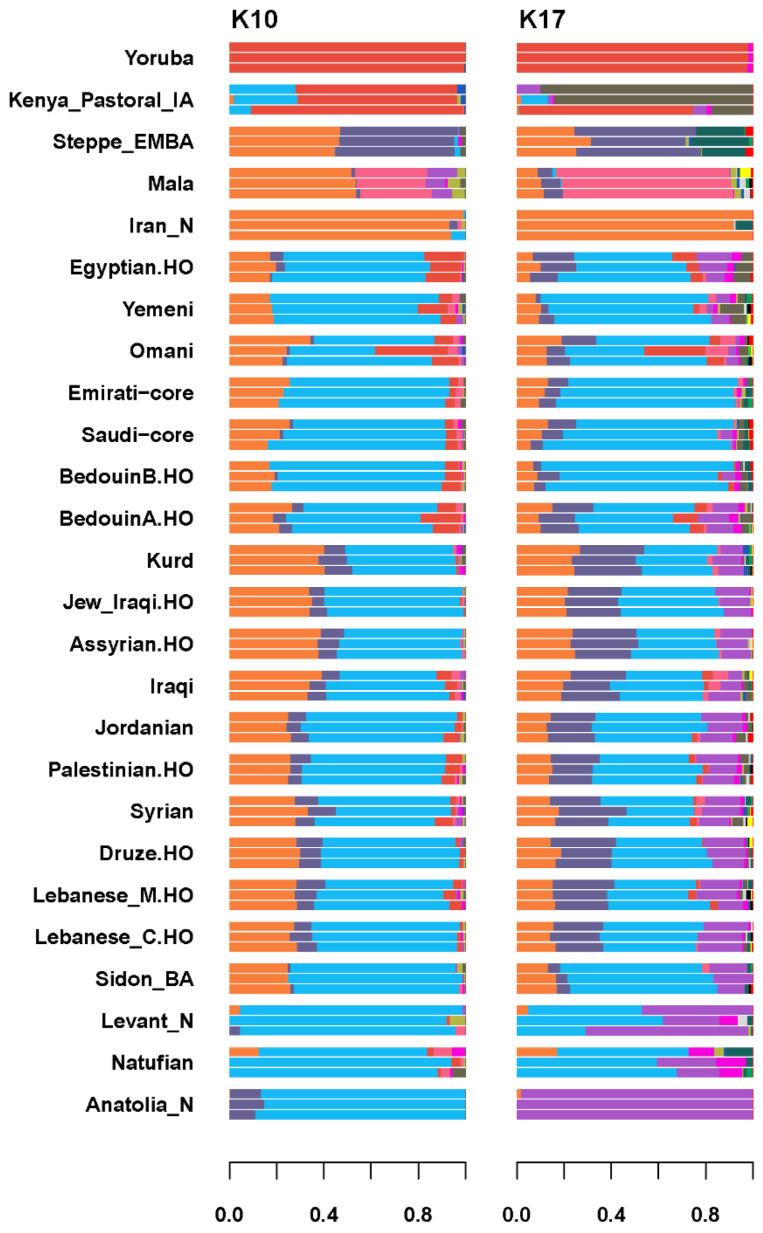


**Figure S6: ADMIXTURE analysis.** Plots using ~80,000 transversions in an unsupervised run and showing results from K = 10 (lowest CV error) and K=17 (differentiation of the Anatolia_N and Natufian components).

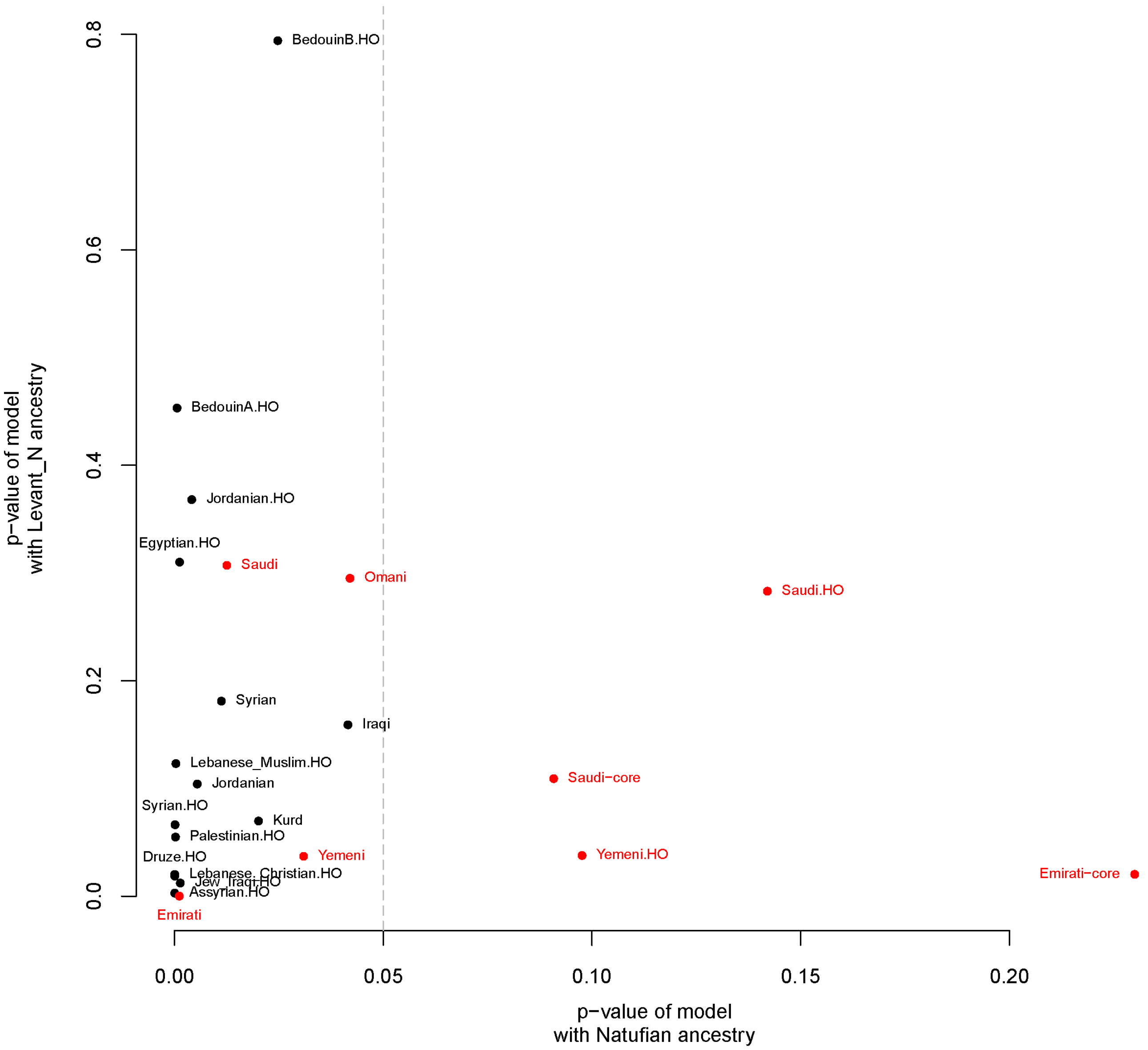


**Figure S7: Testing Levant_N and Natufians as a local source of ancestry for the present-day Middle Easterners.** We plot the p-values from Table S1 and highlight in red the Arabian populations. Saudis, Emiratis, and Yemenis can be modelled (p-value >0.05) with a local ancestry that is Natufian-like without requiring additional ancestry found in Levant_N.

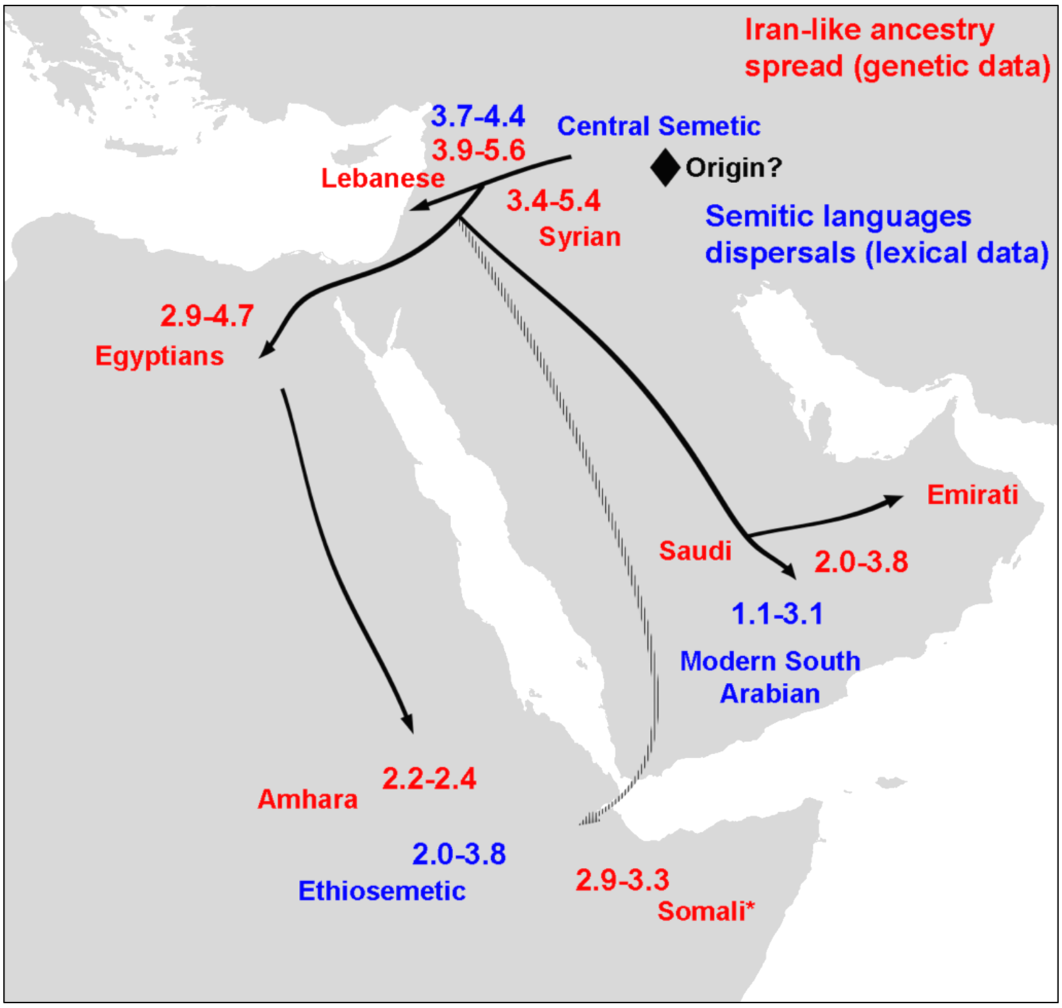


**Figure S8: Spread of Iran-like ancestry and Semitic languages.** Map shows admixture dates in thousands of years ago (red) based on Table S3 and Semitic languages dispersals estimated by Kitchen et al. 2009 from lexical data. Kitchen et al. estimate an Early Bronze Age origin for Semitic ~5.7 KYA in the Levant, and propose that Ethiosemitic was introduced from southern Arabia (dashed arrow) approximately 2.8 KYA. Our admixture tests Table S4 and S5 suggest an Egyptian source of ancestry in East Africa, though ancient DNA from Arabia is still missing to make a comparable analysis. Admixture also appears in non-Semitic speaking populations such as the Somalis.

**A**

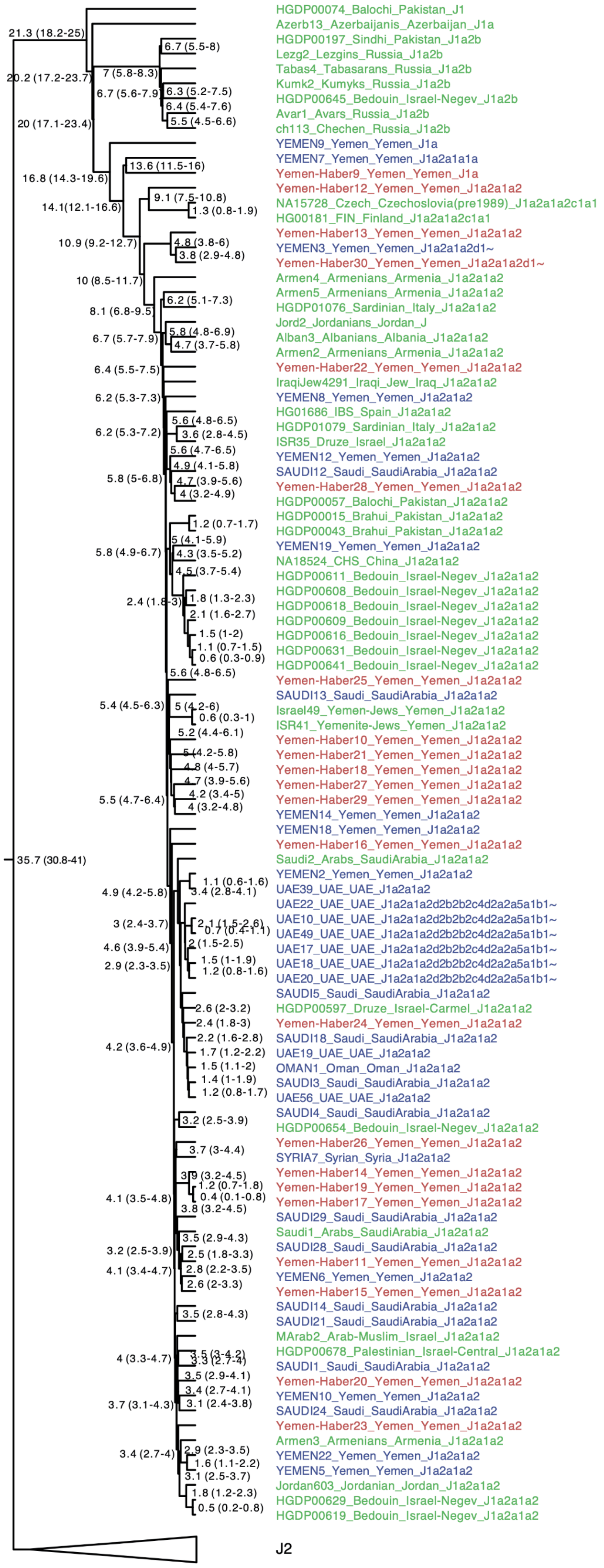
HGDP00074_Balochi_Pakistan_J1
Azerb13_Azerbaijanis_Azerbaijan_J1a
HGDP00197_Sindhi_Pakistan_J1a2b
Lezg2_Lezgins_Russia_J1a2b
Tabas4_Tabasarans_Russia_J1a2b
Kumk2_Kumyks_Russia_J1a2b
HGDP00645_Bedouin_Israel-Negev_J1a2b
Avar1_Avars_Russia_J1a2b
ch113_Chechen_Russia_J1a2b
YEMEN9_Yemen_Yemen_J1a
YEMEN7_Yemen_Yemen_J1a2a1a1a
Yemen-Haber9_Yemen_Yemen_J1a
Yemen-Haber12_Yemen_Yemen_J1a2a1a2
NA15728_Czech_Czechoslovia(pre1989)_J1a2a1a2c1a1
HG00181_FIN_Finland_J1a2a1a2c1a1
Yemen-Haber13_Yemen_Yemen_J1a2a1a2
YEMEN3_Yemen_Yemen_J1a2a1a2d1~
Yemen-Haber30_Yemen_Yemen_J1a2a1a2d1~
Armen4_Armenians_Armenia_J1a2a1a2
Armen5_Armenians_Armenia_J1a2a1a2
HGDP01076_Sardinian_Italy_J1a2a1a2
Jord2_Jordanians_Jordan_J
Alban3_Albanians_Albania_J1a2a1a2
Armen2_Armenians_Armenia_J1a2a1a2
Yemen-Haber22_Yemen_Yemen_J1a2a1a2
IraqiJew4291_Iraqi_Jew_Iraq_J1a2a1a2
YEMEN8_Yemen_Yemen_J1a2a1a2
HG01686_IBS_Spain_J1a2a1a2
HGDP01079_Sardinian_Italy_J1a2a1a2
ISR35_Druze_Israel_J1a2a1a2
YEMEN12_Yemen_Yemen_J1a2a1a2
SAUDI12_Saudi_SaudiArabia_J1a2a1a2
Yemen-Haber28_Yemen_Yemen_J1a2a1a2
HGDP00057_Balochi_Pakistan_J1a2a1a2
HGDP00015_Brahui_Pakistan_J1a2a1a2
HGDP00043_Brahui_Pakistan_J1a2a1a2
YEMEN19_Yemen_Yemen_J1a2a1a2
NA18524_CHS_China_J1a2a1a2
HGDP00611_Bedouin_Israel-Negev_J1a2a1a2
HGDP00608_Bedouin_Israel-Negev_J1a2a1a2
HGDP00618_Bedouin_Israel-Negev_J1a2a1a2
HGDP00609_Bedouin_Israel-Negev_J1a2a1a2
HGDP00616_Bedouin_Israel-Negev_J1a2a1a2
HGDP00631_Bedouin_Israel-Negev_J1a2a1a2
HGDP00641_Bedouin_Israel-Negev_J1a2a1a2
Yemen-Haber25_Yemen_Yemen_J1a2a1a2
SAUDI13_Saudi_SaudiArabia_J1a2a1a2
Israel49_Yemen-Jews_Yemen_J1a2a1a2
ISR41_Yemenite-Jews_Yemen_J1a2a1a2
Yemen-Haber10_Yemen_Yemen_J1a2a1a2
Yemen-Haber21_Yemen_Yemen_J1a2a1a2
Yemen-Haber18_Yemen_Yemen_J1a2a1a2
Yemen-Haber27_Yemen_Yemen_J1a2a1a2
Yemen-Haber29_Yemen_Yemen_J1a2a1a2
YEMEN14_Yemen_Yemen_J1a2a1a2
YEMEN18_Yemen_Yemen_J1a2a1a2
Yemen-Haber16_Yemen_Yemen_J1a2a1a2
Saudi2_Arabs_SaudiArabia_J1a2a1a2
YEMEN2_Yemen_Yemen_J1a2a1a2
UAE39_UAE_UAE_J1a2a1a2
UAE22_UAE_UAE_J1a2a1a2d2b2b2c4d2a2a5a1b1~
UAE10_UAE_UAE_J1a2a1a2d2b2b2c4d2a2a5a1b1~
UAE49_UAE_UAE_J1a2a1a2d2b2b2c4d2a2a5a1b1~
UAE17_UAE_UAE_J1a2a1a2d2b2b2c4d2a2a5a1b1~
UAE18_UAE_UAE_J1a2a1a2d2b2b2c4d2a2a5a1b1~
UAE20_UAE_UAE_J1a2a1a2d2b2b2c4d2a2a5a1b1~
SAUDI5_Saudi_SaudiArabia_J1a2a1a2
HGDP00597_Druze_Israel-Carmel_J1a2a1a2
Yemen-Haber24_Yemen_Yemen_J1a2a1a2
SAUDI18_Saudi_SaudiArabia_J1a2a1a2
UAE19_UAE_UAE_J1a2a1a2
OMAN1_Oman_Oman_J1a2a1a2
SAUDI3_Saudi_SaudiArabia_J1a2a1a2
UAE56_UAE_UAE_J1a2a1a2
SAUDI4_Saudi_SaudiArabia_J1a2a1a2
HGDP00654_Bedouin_Israel-Negev_J1a2a1a2
Yemen-Haber26_Yemen_Yemen_J1a2a1a2
SYRIA7_Syrian_Syria_J1a2a1a2
Yemen-Haber14_Yemen_Yemen_J1a2a1a2
Yemen-Haber19_Yemen_Yemen_J1a2a1a2
Yemen-Haber17_Yemen_Yemen_J1a2a1a2
SAUDI29_Saudi_SaudiArabia_J1a2a1a2
Saudi1_Arabs_SaudiArabia_J1a2a1a2
SAUDI28_Saudi_SaudiArabia_J1a2a1a2
Yemen-Haber11_Yemen_Yemen_J1a2a1a2
YEMEN6_Yemen_Yemen_J1a2a1a2
Yemen-Haber15_Yemen_Yemen_J1a2a1a2
SAUDI14_Saudi_SaudiArabia_J1a2a1a2
SAUDI21_Saudi_SaudiArabia_J1a2a1a2
MArab2_Arab-Muslim_Israel_J1a2a1a2
HGDP00678_Palestinian_Israel-Central_J1a2a1a2
SAUDI1_Saudi_SaudiArabia_J1a2a1a2
Yemen-Haber20_Yemen_Yemen_J1a2a1a2
YEMEN10_Yemen_Yemen_J1a2a1a2
SAUDI24_Saudi_SaudiArabia_J1a2a1a2
Yemen-Haber23_Yemen_Yemen_J1a2a1a2
Armen3_Armenians_Armenia_J1a2a1a2
YEMEN22_Yemen_Yemen_J1a2a1a2
YEMEN5_Yemen_Yemen_J1a2a1a2
Jordan603_Jordanian_Jordan_J1a2a1a2
HGDP00629_Bedouin_Israel-Negev_J1a2a1a2
HGDP00619_Bedouin_Israel-Negev_J1a2a1a2
J2
35.7 (30.8-41)
21.3 (18.2-25)
20.2 (17.2-23.7)
20 (17.1-23.4)
16.8 (14.3-19.6)
14.1(12.1-16.6)
13.6 (11.5-16)
10.9 (9.2-12.7)
10 (8.5-11.7)
9.1 (7.5-10.8)
1.3 (0.8-1.9)
8.1 (6.8-9.5)
7 (5.8-8.3)
6.7 (5.5-8)
6.7 (5.6-7.9)
6.3 (5.2-7.5)
6.4 (5.4-7.6)
5.5 (4.5-6.6)
4.8 (3.8-6)
3.8 (2.9-4.8)
6.2 (5.1-7.3)
6.7 (5.7-7.9)
5.8 (4.8-6.9)
4.7 (3.7-5.8)
6.4 (5.5-7.5)
6.2 (5.3-7.3)
6.2 (5.3-7.2)
5.6 (4.8-6.5)
3.6 (2.8-4.5)
5.6 (4.7-6.5)
4.9 (4.1-5.8)
5.8 (5-6.8)
4.7 (3.9-5.6)
4 (3.2-4.9)
1.2 (0.7-1.7)
5 (4.1-5.9)
5.8 (4.9-6.7)
4.3 (3.5-5.2)
4.5 (3.7-5.4)
2.4 (1.8-3)
1.8 (1.3-2.3)
2.1 (1.6-2.7)
1.5 (1-2)
1.1 (0.7-1.5)
0.6 (0.3-0.9)
5.6 (4.8-6.5)
5.4 (4.5-6.3)
5 (4.2-6)
0.6 (0.3-1)
5.2 (4.4-6.1)
5 (4.2-5.8)
4.8 (4-5.7)
4.7 (3.9-5.6)
5.5 (4.7-6.4)
4.2 (3.4-5)
4 (3.2-4.8)
1.1 (0.6-1.6)
4.9 (4.2-5.8)
3.4 (2.8-4.1)
3 (2.4-3.7)
2.1 (1.5-2.6)
0.7 (0.4-1.1)
4.6 (3.9-5.4)
2 (1.5-2.5)
2.9 (2.3-3.5)
1.5 (1-1.9)
1.2 (0.8-1.6)
2.6 (2-3.2)
2.4 (1.8-3)
2.2 (1.6-2.8)
4.2 (3.6-4.9)
1.7 (1.2-2.2)
1.5 (1.1-2)
1.4 (1-1.9)
1.2 (0.8-1.7)
3.2 (2.5-3.9)
3.7 (3-4.4)
3.9 (3.2-4.5)
1.2 (0.7-1.8)
4.1 (3.5-4.8)
0.4 (0.1-0.8)
3.8 (3.2-4.5)
3.5 (2.9-4.3)
3.2 (2.5-3.9)
2.5 (1.8-3.3)
4.1 (3.4-4.7)
2.8 (2.2-3.5)
2.6 (2-3.3)
3.5 (2.8-4.3)
3.5 (3-4.2)
4 (3.3-4.7)
3.3 (2.7-4)
3.5 (2.9-4.1)
3.4 (2.7-4.1)
3.7 (3.1-4.3)
3.1 (2.4-3.8)
2.9 (2.3-3.5)
3.4 (2.7-4)
1.6 (1.1-2.2)
3.1 (2.5-3.7)
1.8 (1.2-2.3)
0.5 (0.2-0.8)

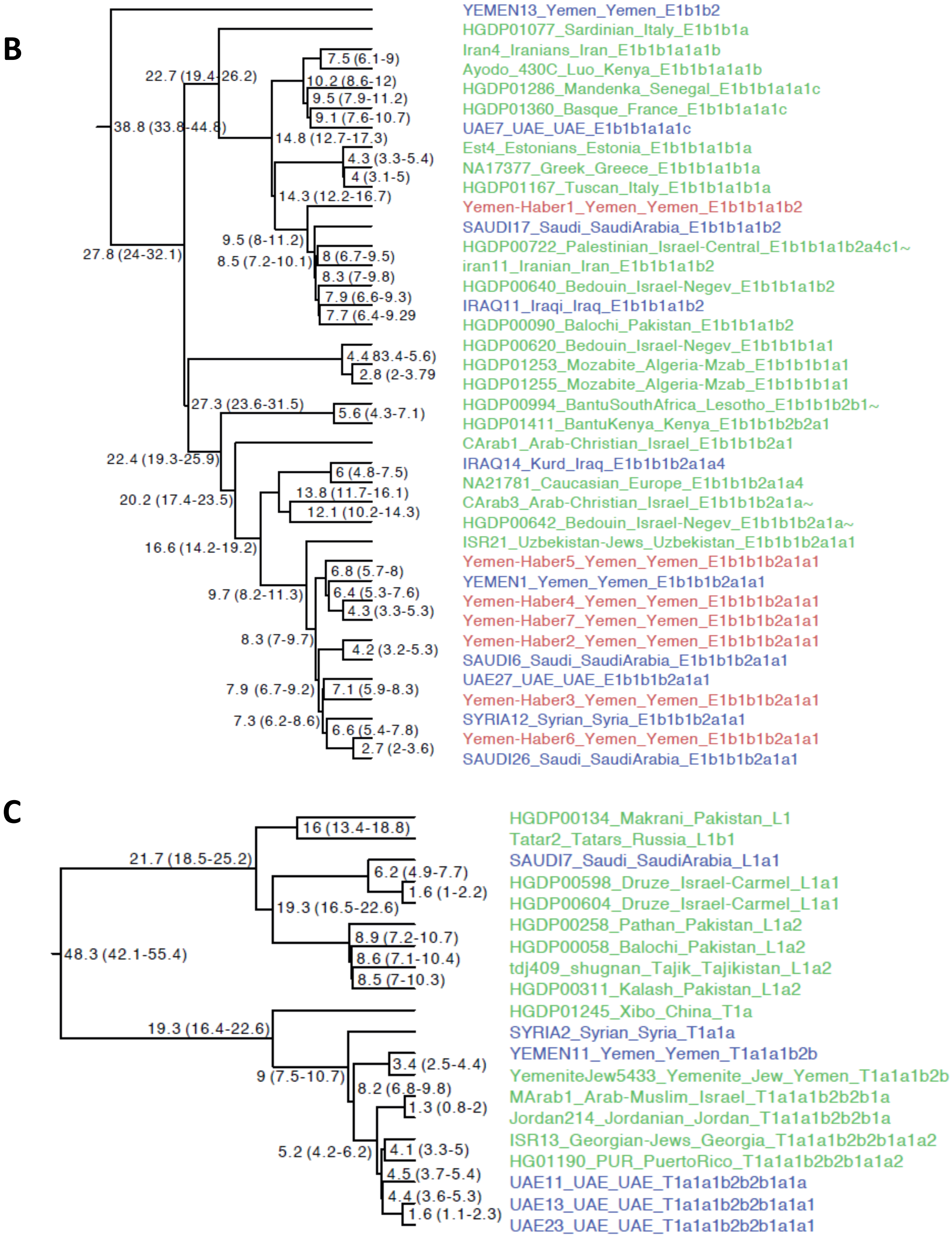


**Figure S9: Y-chromosome phylogeny.** We merged our dataset (samples in Blue) with Haber et al., 2019 (samples in Red) and Hallast et al., 2020 (Samples in Green). We display common haplogroups found in our dataset **(A)** J1, **(B)** E1b and **(C)** L-T. Numbers at each node represent coalescence date in thousand years with 95% confidence intervals in brackets.

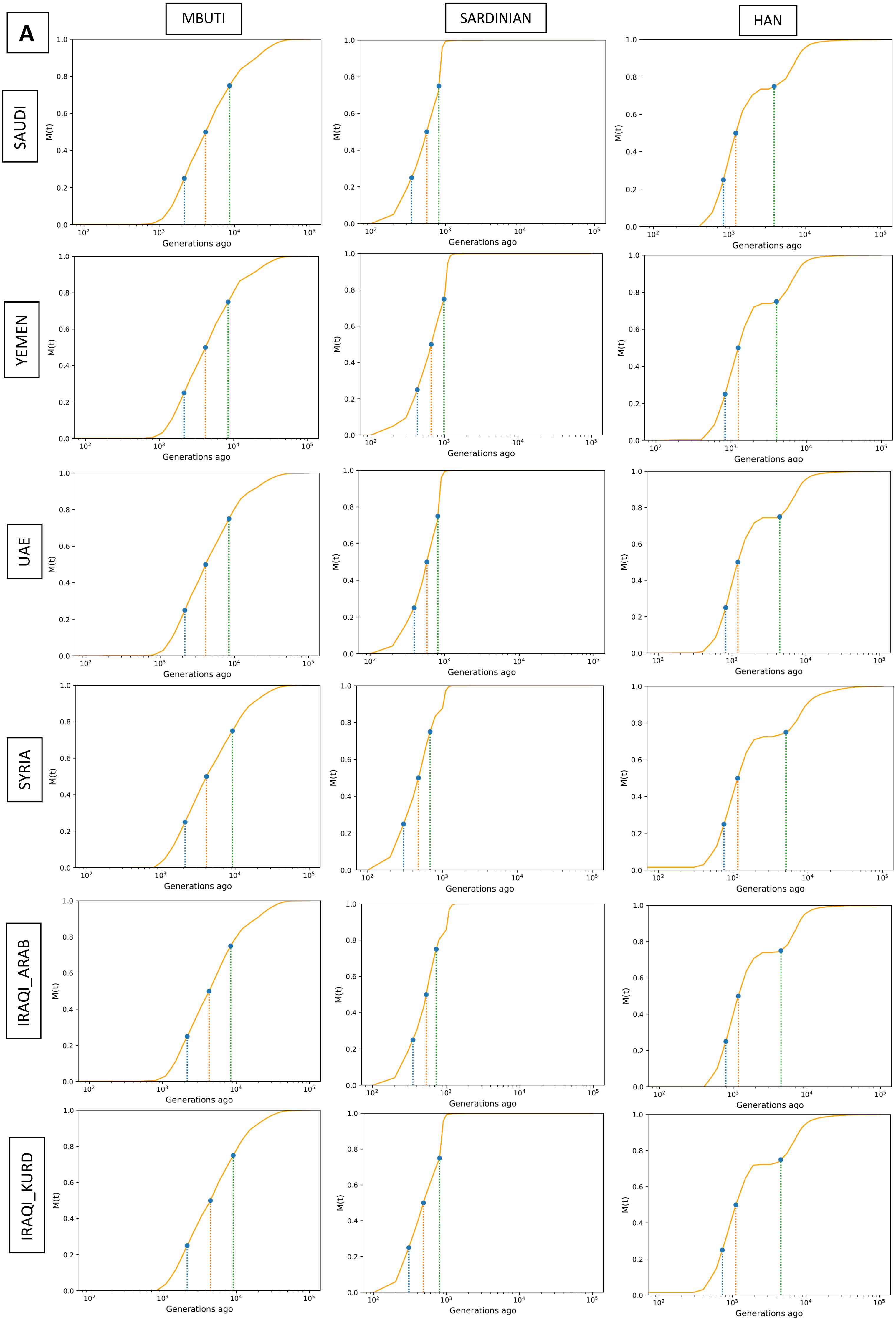

A
MBUTI
SARDINIAN
HAN
SAUDI
YEMEN
UAE
SYRIA
IRAQI_ARAB
IRAQI_KURD
M(t)
Generations ago

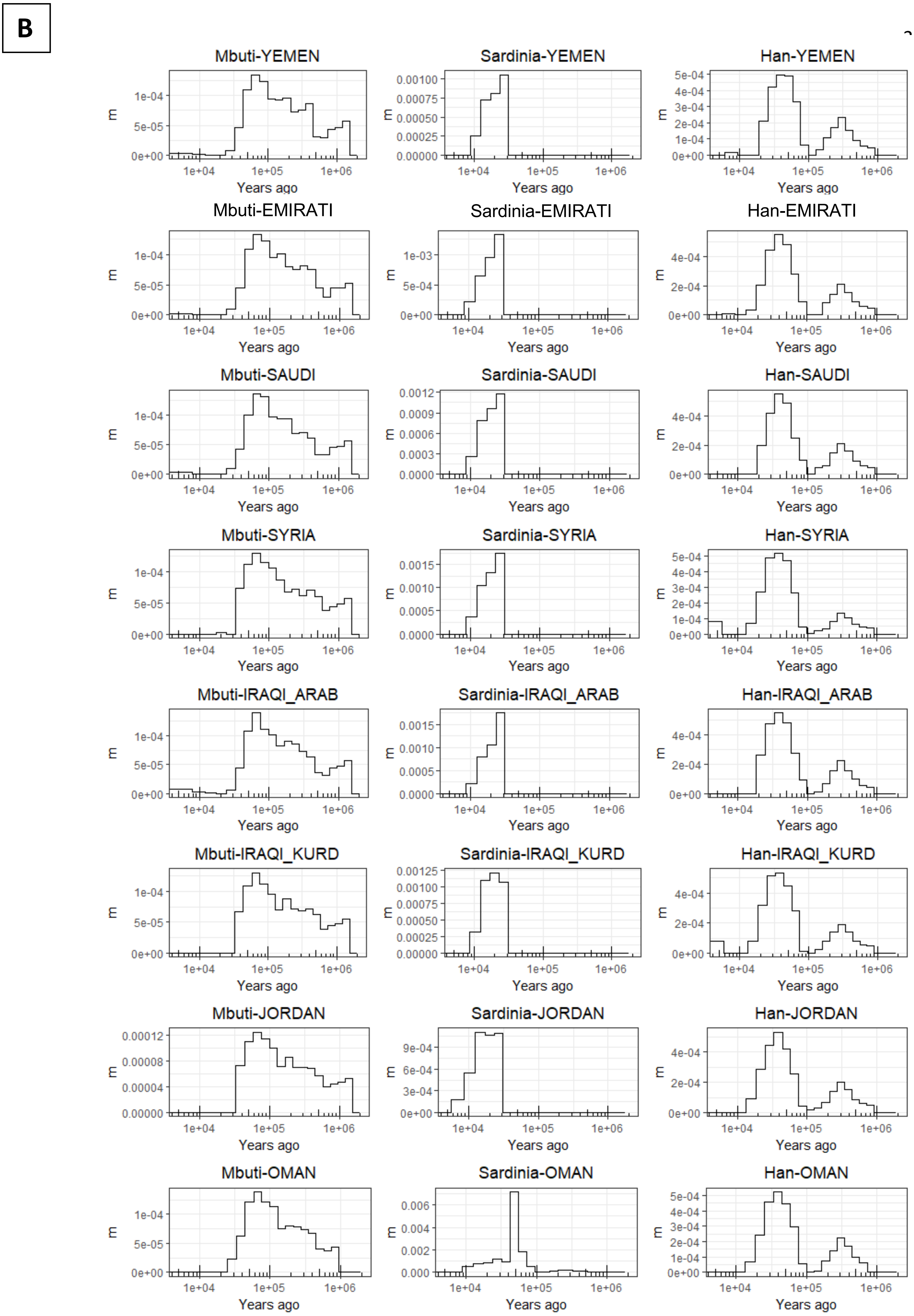


**Figure S10: Migration rates inferred using MSMC-IM**. **A)** Cumulative migration probability, M(t), of Middle Eastern samples compared to Mbuti, Sardinians and Han. Shaded lines illustrate when the M(t) reaches, 25%, 50% and 75%. **B)** Migration rates, m, for the same populations. Note the gradual separation from Mbuti, more of a clean split from Sardinians and the second, older, peak found in the Han comparisons which are consistent with archaic hominin lineages.

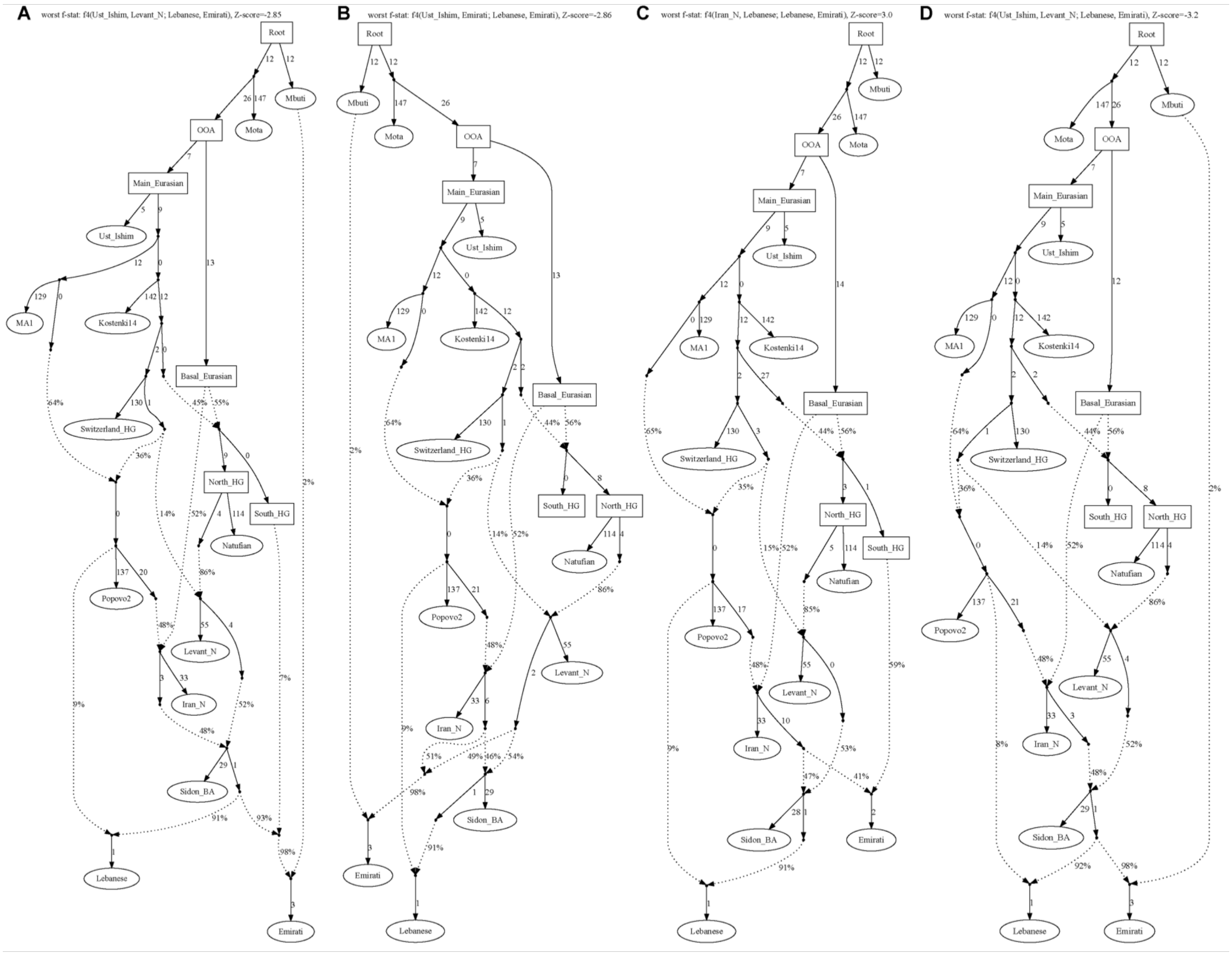


**Figure S11: qpGraph alternative models for population formation in the Middle East.** Graphs show alternative scenarios for populating the Middle East. Changes from the best model (Figure 4) involve (A) Arabians derive their ancestry from a population more closely related to Bronze Age Levantines. (B) Ancestry in Arabia from a Levant_N-related rather than Natufian-related population. (C) A model without additional African ancestry in Arabia. (D) A model without additional Natufian-like ancestry in Arabia. Note the worst f4-statistics on the top of each figure.

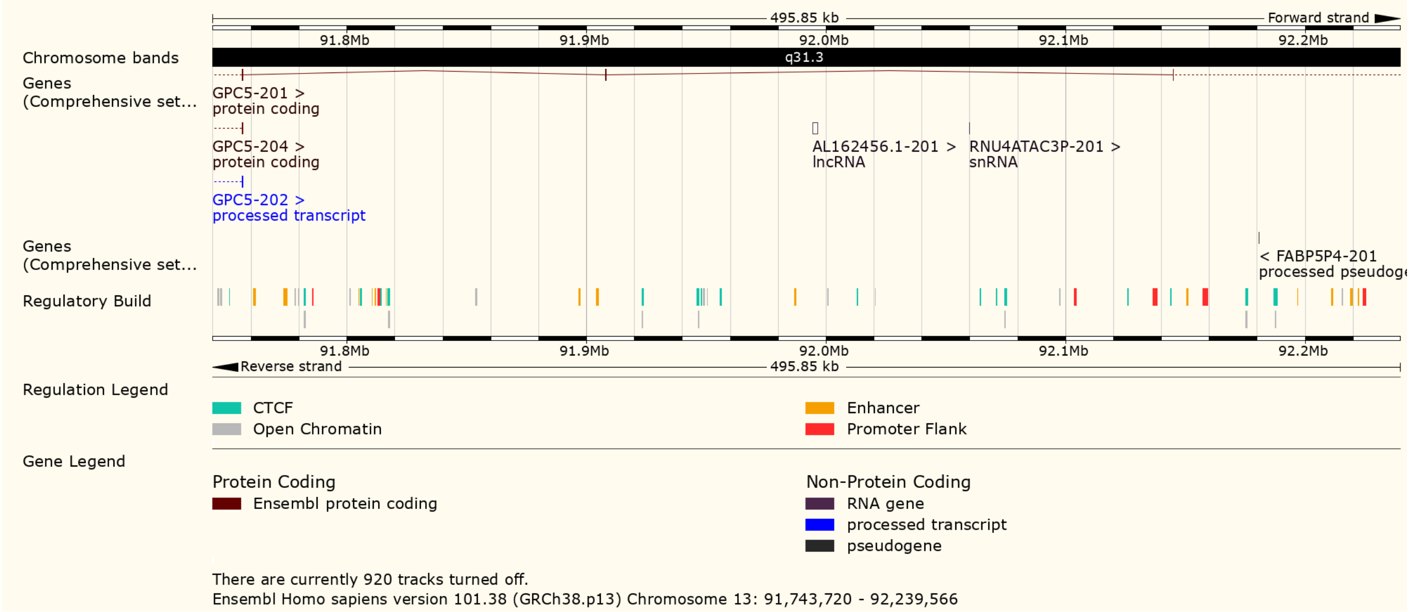


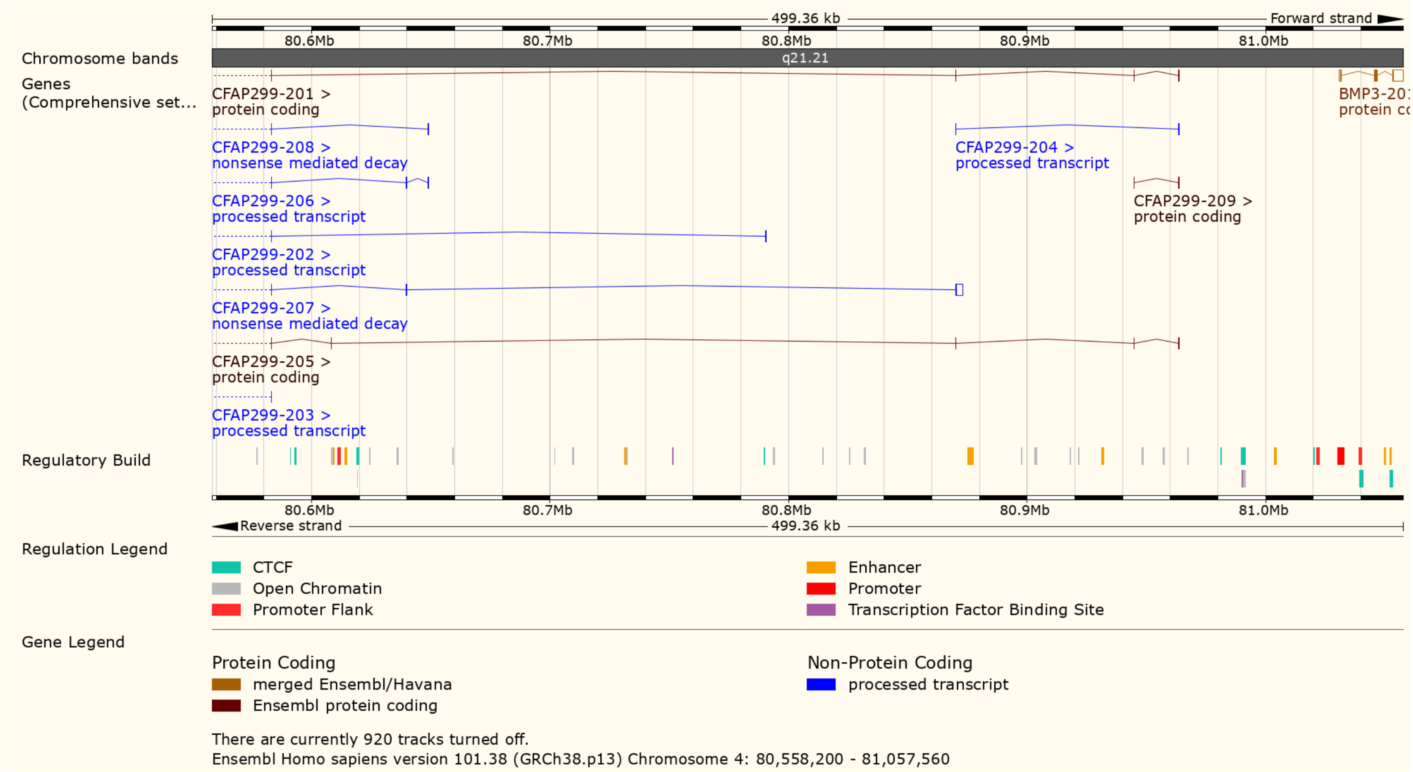


**Figure S12: Neanderthal introgressed segments common in Arabia but rare globally identified using Spime.** Top: 496kb segment on chromosome 13 present at ~20% frequency in Saudi populations but rare globally (Global 1000G Project = 0.02%) and overlapping *GPC5*, a gene expressed in brain tissues. Bottom: 499kb segment on chromosome 4 that reaches ~20% frequency in Emirati.core and overlaps *CFAP299* expressed in the testes with a role in spermatogenesis, and *BMP3*, a cytokine which induces cartilage and bone development (Global 1000G Project < 0.05%). We searched for functional variants within these haplotypes but did not find any amino acid changes within canonical transcripts, with most substitutions limited to introns. Figures downloaded from Ensembl.

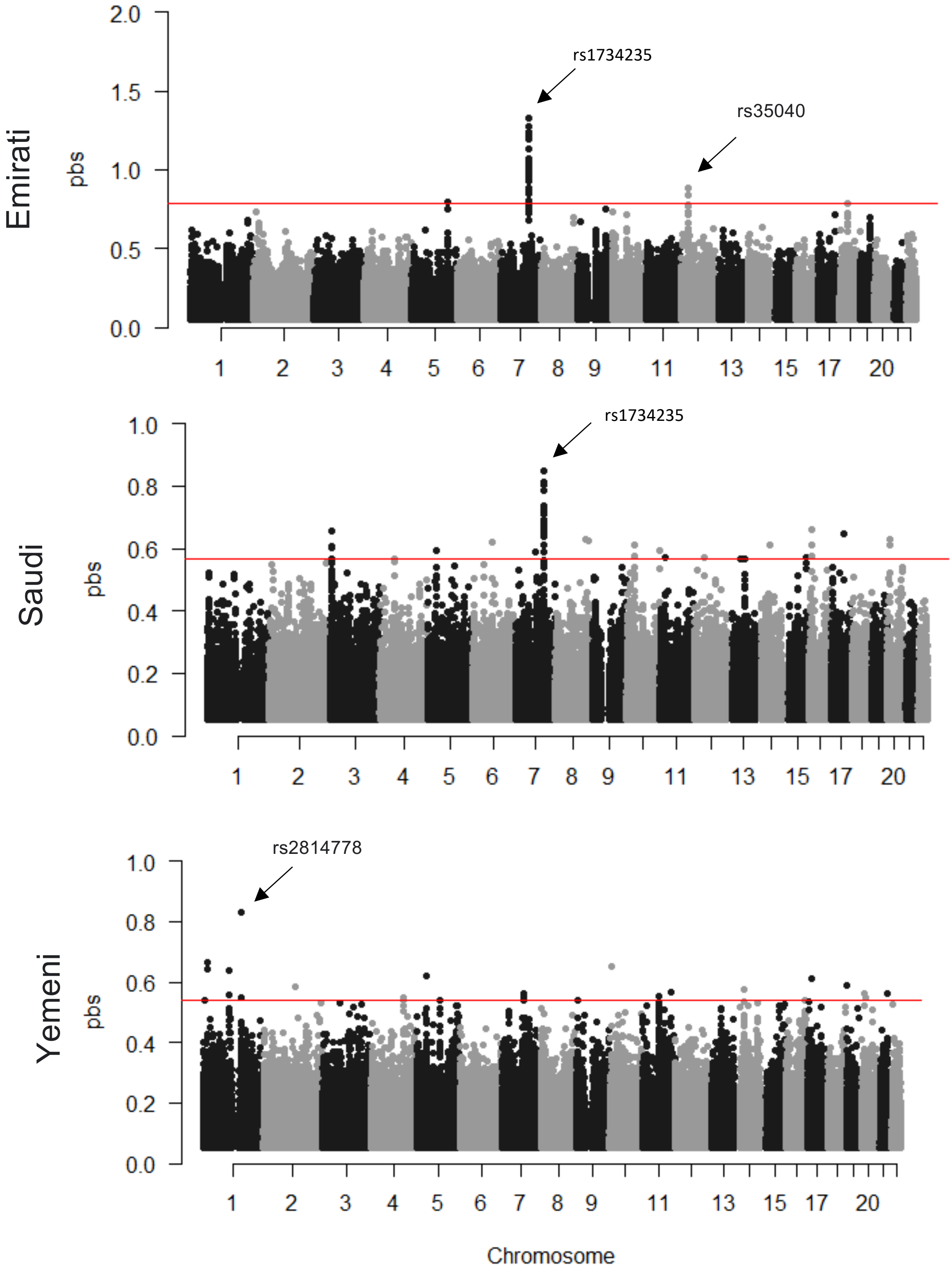


**Figure S13: Population branch statistics (pbs) comparing each Arabian population with Iraqi_Arabs and using Syrians as an outgroup.** Variants showing extreme branch statistics highlighted. Red line illustrates 99.999% quantile. Note the different y-axis scales. rs2814778 is the variant discussed in the main text found at high frequencies in Yemenis that results in the Duffy-null genotype. rs35040 shows strong differentiation in Emiratis and is an eQTL for *DDX11* in multiple tissues. For both Emiratis and Saudis, we find a strong signal of differentiation at a 97kb haplotype on chromosome 7. Variants on this haplotype (rs1734235) almost reaches fixation (97% and 85%, in Emiratis and Saudis respectively) and are associated with increased expression of the lincRNA (AC003088.1).

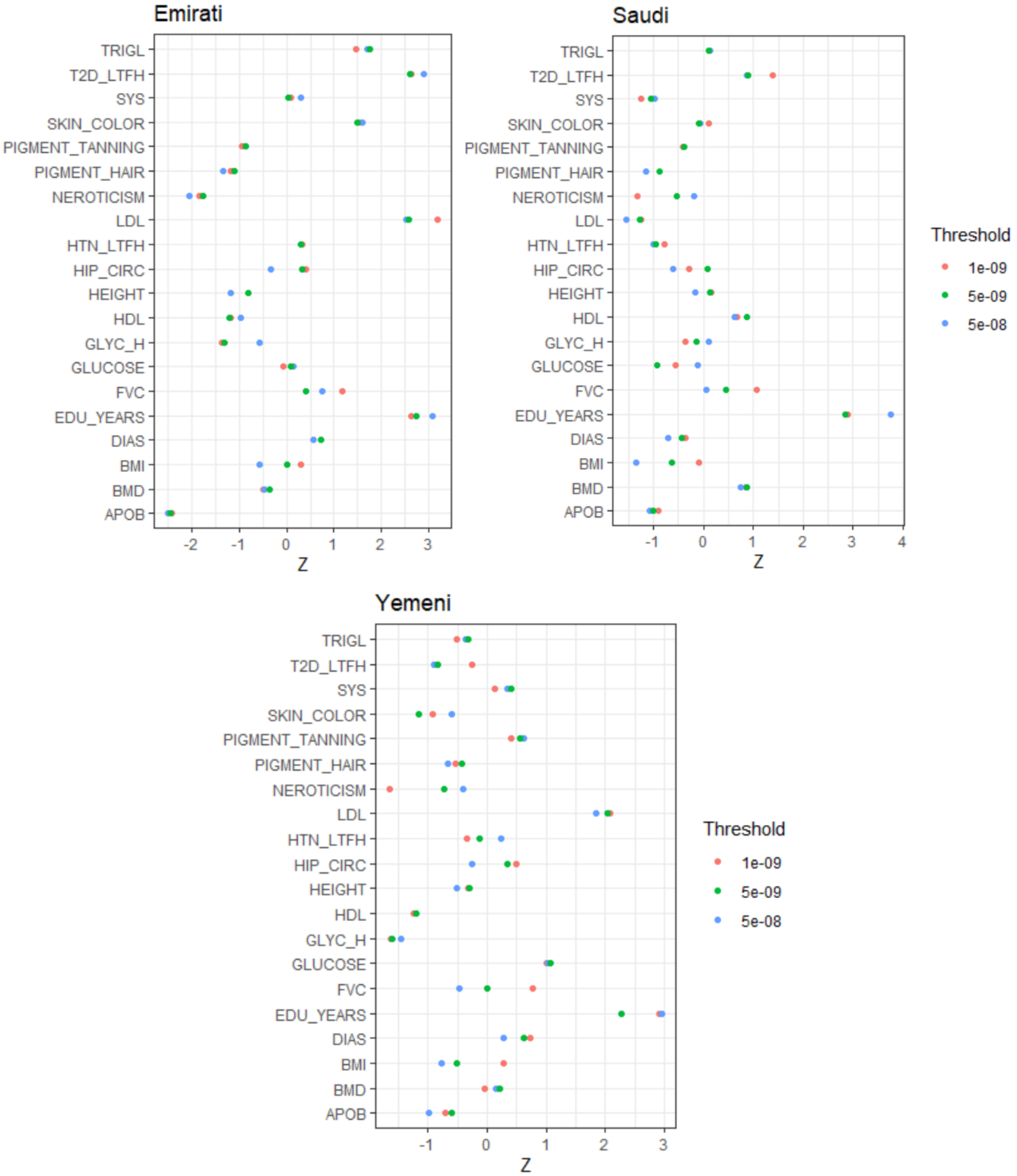


**Figure S14: Effect of significance thresholds on signals of polygenic selection.** For each population, we repeat the test performed in Figure 5 but choosing a more significant p-value threshold (1e-8 and 5e-9). Blocks containing variants not reaching the significant threshold will be dropped from each corresponding analysis. We find very similar values in all significant thresholds. Note the different X-axis scales. LTFH: Association calculated using a liability threshold model, conditional on both case-control status and family history.

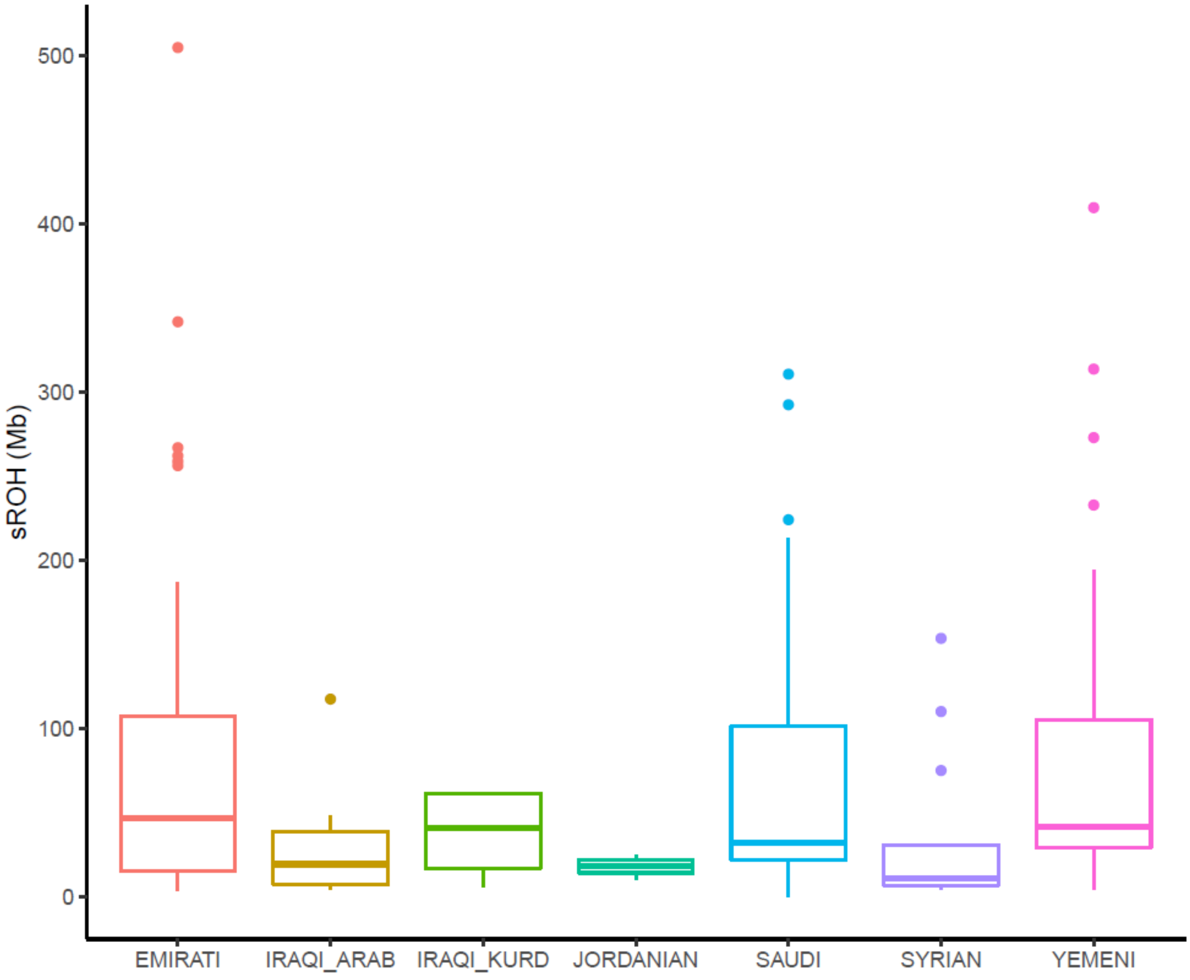


**Figure S15: Sum of runs of homozogysity (sROH) in Megabases (Mb) of samples in this dataset.** sROH using a minimum ROH block of 1Mb (see Methods). The lower and upper hinges in each boxplot correspond to the first and third quartiles (the 25th and 75th percentiles), whisker extends from the hinge to the largest or lowest value no further than 1.5 * Interquartile range. Horizontal line within each boxplot represents the median. Arabians show higher sROH in comparison to Levantines and Iraqis.

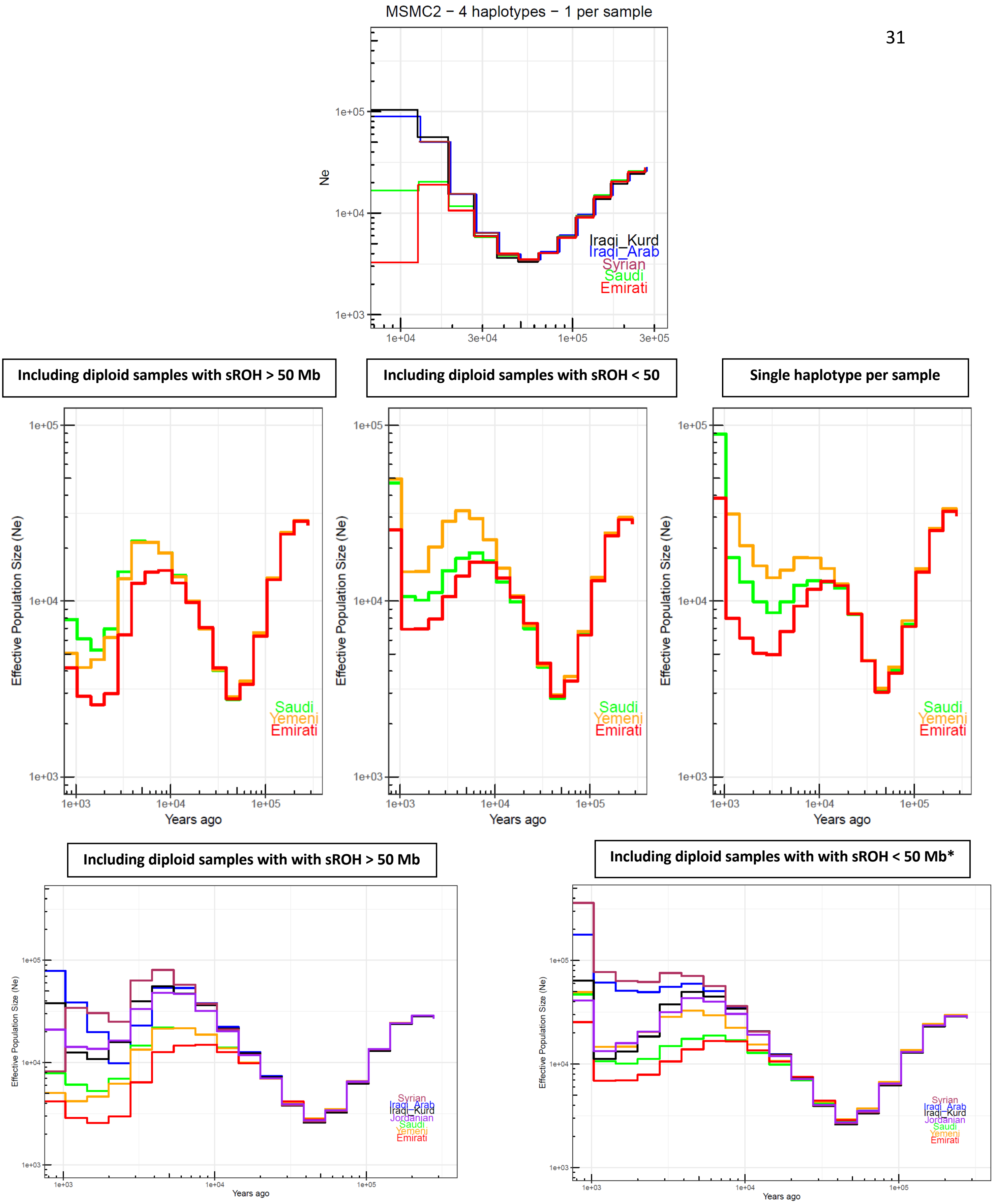


**Figure S16A: Effective population size estimates. Top:** Replicating the divergence in population size between the Levant and Arabia using MSMC2. **Center**: Testing the effect of consanguinity on Arabian population size estimates using Relate**.** sROH calculated using a minimum ROH block of 1Mb. Including samples with likely recent consanguinity affects populations size estimates at recent times. Using a single haplotype per sample reduces this effect. The second bottleneck is apparent in all tests. **Bottom:** Similar to the previous comparisons but including all Middle Eastern populations. Including samples with low sROH show similar results to using a single haplotype per sample show in Figure 2B. *Note that 2 Iraqi-Kurdish samples with ~61Mb sROH were included in this analysis as we have a low number of samples from this population.

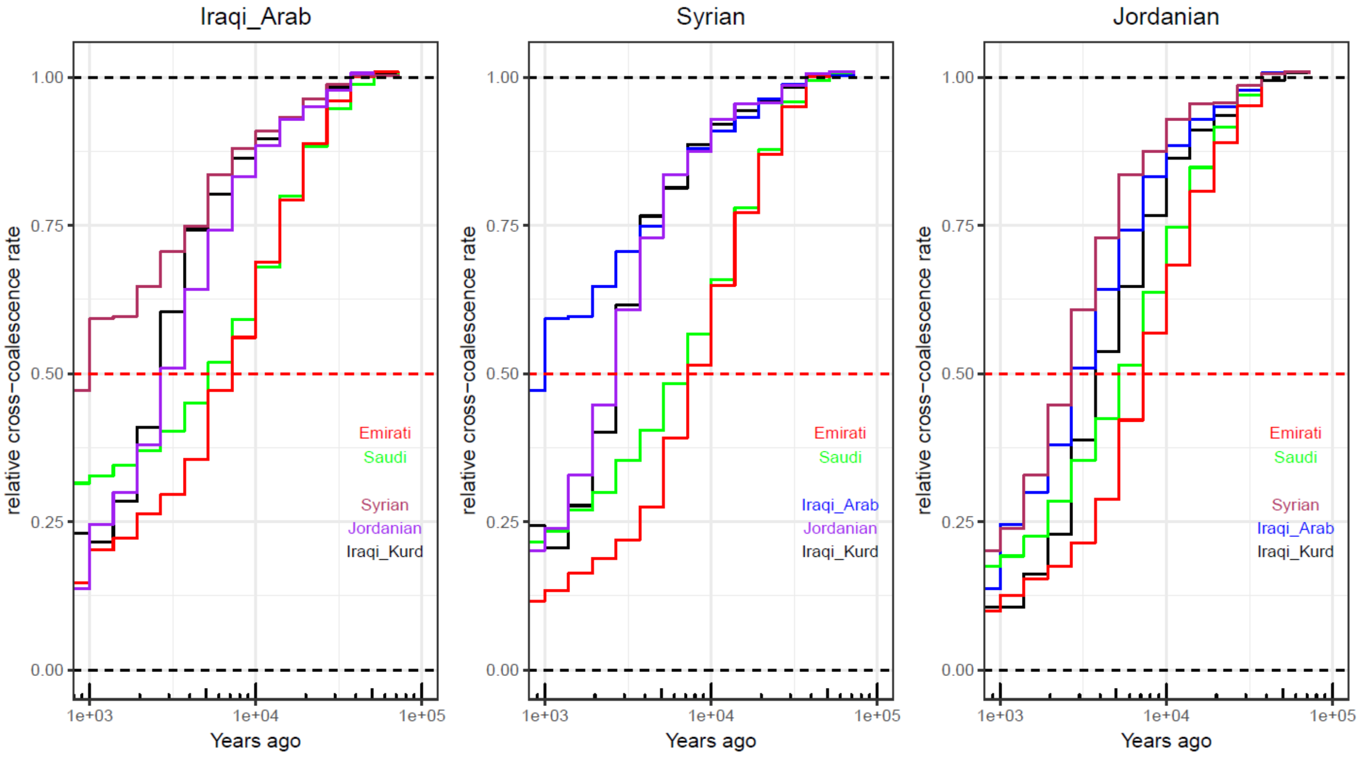


**Figure S16B: Separation History within the Middle East**.
Population indicated at the top of each panel, and within each panel.

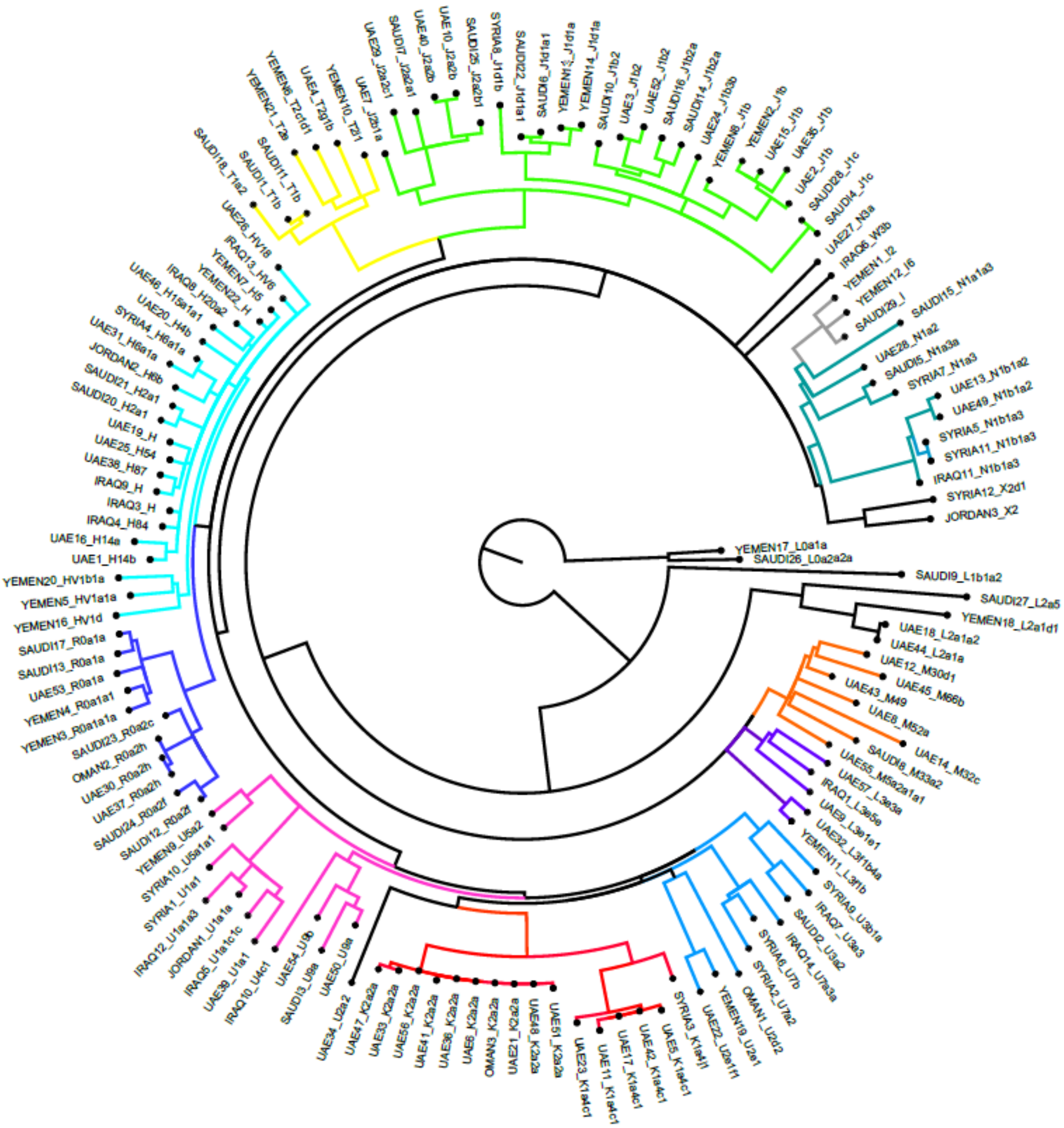


**Figure S17: Maximum-likelihood phylogenetic tree of mitochondrial sequences from samples analysed in this study.** Mitochondrial haplogroup is displayed with sample ID at each node. Similar haplogroups are coloured with the same colour. Branch lengths are not to scale. The effect of African admixture in Arabian populations is observed with L0 haplogroups.

| Test | P value for rank=3 | Ancestry proportions | | | | | | | | P value for rank=3 | Ancestry proportions | | | | | | | |
|---|---|---|---|---|---|---|---|---|---|---|---|---|---|---|---|---|---|---|
| | | Levant_N | SE | Iran_N | SE | EHG | SE | Mota | SE | | Natufian | SE | Iran_N | SE | EHG | SE | Mota | SE |
| Assyrian.HO | 2.78E-03 | 0.32 | 0.02 | 0.61 | 0.02 | 0.10 | 0.01 | -0.02 | 0.01 | 8.00E-06 | 0.38 | 0.02 | 0.56 | 0.02 | 0.09 | 0.01 | -0.03 | 0.01 |
| BedouinA.HO | **4.53E-01** | 0.42 | 0.02 | 0.39 | 0.02 | 0.09 | 0.01 | 0.09 | 0.01 | 5.70E-04 | 0.48 | 0.02 | 0.36 | 0.02 | 0.09 | 0.01 | 0.08 | 0.01 |
| BedouinB.HO | **7.94E-01** | 0.54 | 0.02 | 0.35 | 0.02 | 0.06 | 0.02 | 0.05 | 0.01 | 2.47E-02 | 0.56 | 0.02 | 0.32 | 0.02 | 0.07 | 0.01 | 0.04 | 0.01 |
| Druze.HO | 1.86E-02 | 0.39 | 0.02 | 0.49 | 0.02 | 0.13 | 0.01 | 0.00 | 0.01 | 5.00E-06 | 0.45 | 0.02 | 0.44 | 0.02 | 0.12 | 0.01 | -0.01 | 0.01 |
| Egyptian.HO | **3.10E-01** | 0.45 | 0.02 | 0.32 | 0.02 | 0.08 | 0.01 | 0.15 | 0.01 | 1.18E-03 | 0.50 | 0.02 | 0.30 | 0.02 | 0.08 | 0.01 | 0.12 | 0.01 |
| Iraqi_Arab | **1.59E-01** | 0.31 | 0.02 | 0.54 | 0.02 | 0.12 | 0.01 | 0.03 | 0.01 | 4.15E-02 | 0.38 | 0.02 | 0.49 | 0.02 | 0.11 | 0.01 | 0.02 | 0.01 |
| Jew_Iraqi.HO | 1.21E-02 | 0.35 | 0.02 | 0.55 | 0.02 | 0.11 | 0.02 | -0.01 | 0.01 | 1.35E-03 | 0.41 | 0.02 | 0.51 | 0.02 | 0.10 | 0.01 | -0.02 | 0.01 |
| Jordanian.HO | **3.68E-01** | 0.38 | 0.02 | 0.43 | 0.02 | 0.13 | 0.01 | 0.06 | 0.01 | 4.09E-03 | 0.46 | 0.02 | 0.38 | 0.02 | 0.11 | 0.01 | 0.05 | 0.01 |
| Jordanian | **1.04E-01** | 0.43 | 0.03 | 0.44 | 0.03 | 0.11 | 0.02 | 0.02 | 0.01 | 5.41E-03 | 0.48 | 0.02 | 0.40 | 0.03 | 0.11 | 0.02 | 0.01 | 0.01 |
| Iraqi_Kurd | **6.97E-02** | 0.23 | 0.02 | 0.62 | 0.02 | 0.16 | 0.02 | -0.01 | 0.01 | 2.01E-02 | 0.31 | 0.02 | 0.56 | 0.03 | 0.15 | 0.02 | -0.01 | 0.01 |
| Lebanese_Christian.HO | 2.01E-02 | 0.42 | 0.02 | 0.46 | 0.02 | 0.13 | 0.01 | -0.01 | 0.01 | 9.00E-06 | 0.49 | 0.02 | 0.41 | 0.02 | 0.12 | 0.01 | -0.02 | 0.01 |
| Lebanese_Muslim.HO | **1.23E-01** | 0.39 | 0.02 | 0.48 | 0.02 | 0.11 | 0.01 | 0.02 | 0.01 | 2.95E-04 | 0.45 | 0.02 | 0.44 | 0.02 | 0.11 | 0.01 | 0.01 | 0.01 |
| Omani | **2.95E-01** | 0.41 | 0.03 | 0.41 | 0.03 | 0.09 | 0.02 | 0.10 | 0.01 | 4.20E-02 | 0.46 | 0.02 | 0.37 | 0.03 | 0.09 | 0.02 | 0.09 | 0.01 |
| Palestinian.HO | **5.48E-02** | 0.40 | 0.02 | 0.43 | 0.02 | 0.11 | 0.01 | 0.06 | 0.01 | 2.02E-04 | 0.47 | 0.02 | 0.39 | 0.02 | 0.10 | 0.01 | 0.04 | 0.01 |
| Saudi.core | **1.09E-01** | 0.49 | 0.02 | 0.42 | 0.02 | 0.06 | 0.01 | 0.03 | 0.01 | **9.08E-02** | 0.52 | 0.02 | 0.39 | 0.02 | 0.07 | 0.01 | 0.02 | 0.01 |
| Saudi | **3.07E-01** | 0.50 | 0.02 | 0.32 | 0.02 | 0.05 | 0.01 | 0.14 | 0.01 | 1.25E-02 | 0.50 | 0.02 | 0.31 | 0.02 | 0.07 | 0.01 | 0.12 | 0.01 |
| Saudi.HO | **2.83E-01** | 0.50 | 0.02 | 0.40 | 0.02 | 0.07 | 0.02 | 0.04 | 0.01 | **1.42E-01** | 0.51 | 0.02 | 0.38 | 0.02 | 0.09 | 0.01 | 0.03 | 0.01 |
| Syrian | **1.81E-01** | 0.34 | 0.02 | 0.50 | 0.02 | 0.14 | 0.01 | 0.02 | 0.01 | 1.12E-02 | 0.40 | 0.02 | 0.46 | 0.02 | 0.13 | 0.01 | 0.01 | 0.01 |
| Syrian.HO | **6.62E-02** | 0.38 | 0.02 | 0.45 | 0.02 | 0.12 | 0.01 | 0.05 | 0.01 | 9.50E-05 | 0.44 | 0.02 | 0.41 | 0.02 | 0.11 | 0.01 | 0.04 | 0.01 |
| Emirati.core | 2.02E-02 | 0.49 | 0.02 | 0.43 | 0.02 | 0.06 | 0.01 | 0.03 | 0.01 | **2.30E-01** | 0.53 | 0.02 | 0.39 | 0.02 | 0.07 | 0.01 | 0.02 | 0.01 |
| Emirati | 2.00E-06 | 0.29 | 0.02 | 0.52 | 0.02 | 0.09 | 0.01 | 0.10 | 0.00 | 1.11E-03 | 0.34 | 0.02 | 0.48 | 0.02 | 0.09 | 0.01 | 0.09 | 0.00 |
| Yemeni | 3.69E-02 | 0.52 | 0.02 | 0.35 | 0.02 | 0.04 | 0.01 | 0.09 | 0.01 | 3.09E-02 | 0.55 | 0.02 | 0.32 | 0.02 | 0.06 | 0.01 | 0.08 | 0.01 |
| Yemeni.HO | 3.77E-02 | 0.38 | 0.02 | 0.40 | 0.02 | 0.06 | 0.01 | 0.16 | 0.01 | **9.76E-02** | 0.42 | 0.02 | 0.37 | 0.02 | 0.07 | 0.01 | 0.14 | 0.01 |

**Table S1.** Modelling present-day Middle Easterners as deriving their ancestry from four ancient populations using qpAdm. We used seven outgroups in the test: Ust'-Ishim, Kostenki14, WHG, CHG, Natufian (or Levant_N), Papuan, and Mbuti. SE = standard error. P value > 0.05 (bold) indicates the model is not rejected. ".HO" - samples from the Human Origins dataset. '.core' represents the curated samples, samples without '.core' represent the general population without the '.core' samples.

| | Reference population | | Admixture | | | |
|---|---|---|---|---|---|---|
| Test | A | B | LD curve amplitude | Z-score | Time (gen ago) | Z-score |
| Saudi.core* | Druze | ITU.SG | 0.00000741567 +/-0.000000411159 | Z=18.036 | 64.2792 +/-11.064 | Z=5.80975 |
| | Druze | PJL.SG | 0.00000380233 +/-0.000000652193 | Z=5.83007 | 64.2792 +/-11.064 | Z=5.80975 |
| | LWK.SG | Druze | 0.0000867775 +/-0.0000145912 | Z=5.94726 | 64.2792 +/-11.064 | Z=5.80975 |
| | LWK.SG | ITU.SG | 0.0000659673 +/-0.0000127224 | Z=5.18513 | 64.2792 +/-11.064 | Z=5.80975 |
| | LWK.SG | Iranian | 0.000079526 +/-0.0000163563 | Z=4.86211 | 64.2792 +/-11.064 | Z=5.80975 |
| | LWK.SG | PJL.SG | 0.0000710359 +/-0.0000133445 | Z=5.32322 | 64.2792 +/-11.064 | Z=5.80975 |
| | Yoruba | Druze | 0.0000915487 +/-0.0000150628 | Z=6.07781 | 64.2792 +/-11.064 | Z=5.80975 |
| | Yoruba | ITU.SG | 0.0000715227 +/-0.0000125884 | Z=5.68163 | 64.2792 +/-11.064 | Z=5.80975 |
| | Yoruba | PJL.SG | 0.0000761204 +/-0.0000139804 | Z=5.44479 | 64.2792 +/-11.064 | Z=5.80975 |
| Emirati.core | LWK.SG | Druze | 0.0000589084 +/-0.00000532686 | Z=11.0588 | 21.5731 +/-2.87294 | Z=7.50907 |
| | LWK.SG | ITU.SG | 0.0000520471 +/-0.00000445911 | Z=11.6721 | 21.5731 +/-2.87294 | Z=7.50907 |
| | LWK.SG | Iranian | 0.000059676 +/-0.00000503894 | Z=11.843 | 21.5731 +/-2.87294 | Z=7.50907 |
| | LWK.SG | PJL.SG | 0.0000526636 +/-0.00000508227 | Z=10.3622 | 21.5731 +/-2.87294 | Z=7.50907 |
| | Yoruba | Druze | 0.0000634949 +/-0.00000577752 | Z=10.99 | 21.5731 +/-2.87294 | Z=7.50907 |
| | Yoruba | ITU.SG | 0.0000555435 +/-0.00000478698 | Z=11.603 | 21.5731 +/-2.87294 | Z=7.50907 |
| | Yoruba | Iranian | 0.0000637986 +/-0.00000540742 | Z=11.7984 | 21.5731 +/-2.87294 | Z=7.50907 |
| | Yoruba | PJL.SG | 0.0000560743 +/-0.00000544697 | Z=10.2946 | 21.5731 +/-2.87294 | Z=7.50907 |
| Iraqi_Arab | Druze | ITU.SG | 0.0000113501 +/-0.00000161324 | Z=7.03563 | 29.0865 +/-6.01517 | Z=4.83553 |
| | Iranian | ITU.SG | 0.0000055752 +/-0.000000684234 | Z=8.14809 | 29.0865 +/-6.01517 | Z=4.83553 |
| | LWK.SG | Druze | 0.0000902094 +/-0.0000131962 | Z=6.83601 | 29.0865 +/-6.01517 | Z=4.83553 |
| | LWK.SG | ITU.SG | 0.0000718344 +/-0.0000103802 | Z=6.92034 | 29.0865 +/-6.01517 | Z=4.83553 |
| | LWK.SG | Iranian | 0.0000894593 +/-0.0000133348 | Z=6.70872 | 29.0865 +/-6.01517 | Z=4.83553 |
| | LWK.SG | PJL.SG | 0.0000759272 +/-0.0000106155 | Z=7.15252 | 29.0865 +/-6.01517 | Z=4.83553 |
| | Yoruba | Druze | 0.0000948199 +/-0.0000140899 | Z=6.72963 | 29.0865 +/-6.01517 | Z=4.83553 |
| | Yoruba | ITU.SG | 0.0000761279 +/-0.000011093 | Z=6.86268 | 29.0865 +/-6.01517 | Z=4.83553 |
| | Yoruba | Iranian | 0.0000936656 +/-0.0000141459 | Z=6.62141 | 29.0865 +/-6.01517 | Z=4.83553 |
| | Yoruba | PJL.SG | 0.0000809727 +/-0.0000114702 | Z=7.05939 | 29.0865 +/-6.01517 | Z=4.83553 |
| Syria | LWK.SG | Druze | 0.000067166 +/-0.00000920866 | Z=7.29378 | 19.1781 +/-4.16062 | Z=4.60945 |
| | LWK.SG | ITU.SG | 0.0000607542 +/-0.00000801921 | Z=7.57609 | 19.1781 +/-4.16062 | Z=4.60945 |
| | LWK.SG | Iranian | 0.0000679894 +/-0.00000883859 | Z=7.69233 | 19.1781 +/-4.16062 | Z=4.60945 |
| | LWK.SG | PJL.SG | 0.0000608678 +/-0.0000083615 | Z=7.27954 | 19.1781 +/-4.16062 | Z=4.60945 |
| | Yoruba | Druze | 0.0000713981 +/-0.0000100742 | Z=7.0872 | 19.1781 +/-4.16062 | Z=4.60945 |
| | Yoruba | ITU.SG | 0.0000646977 +/-0.00000872059 | Z=7.41896 | 19.1781 +/-4.16062 | Z=4.60945 |
| | Yoruba | Iranian | 0.0000724506 +/-0.00000970008 | Z=7.46907 | 19.1781 +/-4.16062 | Z=4.60945 |
| | Yoruba | PJL.SG | 0.0000648224 +/-0.00000906419 | Z=7.15149 | 19.1781 +/-4.16062 | Z=4.60945 |
| Yemen | LWK.SG | Druze | 0.000125983 +/-0.0000116452 | Z=10.8185 | 18.3648 +/-2.59716 | Z=7.0711 |
| | LWK.SG | ITU.SG | 0.0000949767 +/-0.00000903391 | Z=10.5134 | 18.3648 +/-2.59716 | Z=7.0711 |
| | LWK.SG | Iranian | 0.000121224 +/-0.0000112598 | Z=10.7661 | 18.3648 +/-2.59716 | Z=7.0711 |
| | LWK.SG | PJL.SG | 0.000100818 +/-0.00000950151 | Z=10.6107 | 18.3648 +/-2.59716 | Z=7.0711 |
| | Yoruba | Druze | 0.000136766 +/-0.0000123486 | Z=11.0754 | 18.3648 +/-2.59716 | Z=7.0711 |
| | Yoruba | ITU.SG | 0.000104641 +/-0.00000948777 | Z=11.029 | 18.3648 +/-2.59716 | Z=7.0711 |
| | Yoruba | Iranian | 0.000130557 +/-0.0000121187 | Z=10.7732 | 18.3648 +/-2.59716 | Z=7.0711 |
| | Yoruba | PJL.SG | 0.000112707 +/-0.00000995706 | Z=11.3193 | 18.3648 +/-2.59716 | Z=7.0711 |

**Table S2. Testing modern populations for admixture using via Malder**. *Saudi.core have a second recent pulse of admixture identified at 13.7236 +/- 3.63366 generations ago.

| | Reference population | | Admixture | | | |
|---|---|---|---|---|---|---|
| Test | A | B | LD curve amplitude | Z-score | Time (gens ago) | Z-score |
| **Lebanese** | Levant_ChL | Iran_Tepe_Hissar_ChL | 6.61304e-05 +/- 1.56944e-05 | Z=4.21363 | 170.287 +/- 30.9204 | Z=5.50728 |
| | Levant_N | Iran_Hajji_Firuz_ChL | 7.49317e-05 +/- 1.71242e-05 | Z=4.37579 | 170.287 +/- 30.9204 | Z=5.50728 |
| | Levant_N | Iran_N | 0.000132748 +/- 3.28994e-05 | Z=4.03496 | 170.287 +/- 30.9204 | Z=5.50728 |
| | Levant_N | Iran_Seh_Gabi_ChL | 7.70897e-05 +/- 1.88219e-05 | Z=4.09574 | 170.287 +/- 30.9204 | Z=5.50728 |
| | Levant_N | Iran_Tepe_Hissar_ChL | 9.59566e-05 +/- 2.34421e-05 | Z=4.09334 | 170.287 +/- 30.9204 | Z=5.50728 |
| | Natufian | Iran_Hajji_Firuz_ChL | 8.4556e-05 +/- 1.82914e-05 | Z=4.62273 | 170.287 +/- 30.9204 | Z=5.50728 |
| **Syrian*** | Levant_ChL | Iran_N | 0.000111442 +/- 3.04679e-05 | Z=3.65768 | 157.872 +/- 34.498 | Z=4.57628 |
| | Natufian | Iran_Tepe_Hissar_ChL | 8.70245e-05 +/- 2.77751e-05 | Z=3.13318 | 157.872 +/- 34.498 | Z=4.57628 |
| **Arabian.core** | Levant_ChL | Iran_N | 5.99811e-05 +/- 1.88461e-05 | Z=3.18267 | 103.853 +/- 32.7796 | Z=3.16823 |
| | Levant_ChL | Iran_Tepe_Hissar_ChL | 4.64144e-05 +/- 1.4773e-05 | Z=3.14183 | 103.853 +/- 32.7796 | Z=3.16823 |
| | Levant_N | Iran_Tepe_Hissar_ChL | 4.62203e-05 +/- 1.45843e-05 | Z=3.16918 | 103.853 +/- 32.7796 | Z=3.16823 |
| **Egyptian** | Iran_N | Iran_Seh_Gabi_ChL | 2.33472e-05 +/- 6.87091e-06 | Z=3.39798 | 137.114 +/- 31.69 | Z=4.32673 |
| | Levant_ChL | Iran_Hajji_Firuz_ChL | 3.03437e-05 +/- 8.07696e-06 | Z=3.75682 | 137.114 +/- 31.69 | Z=4.32673 |
| | Levant_ChL | Iran_N | 7.8149e-05 +/- 1.93651e-05 | Z=4.03556 | 137.114 +/- 31.69 | Z=4.32673 |
| | Levant_ChL | Iran_Seh_Gabi_ChL | 3.87449e-05 +/- 1.06208e-05 | Z=3.64801 | 137.114 +/- 31.69 | Z=4.32673 |
| | Levant_ChL | Iran_Tepe_Hissar_ChL | 4.32601e-05 +/- 1.11383e-05 | Z=3.88391 | 137.114 +/- 31.69 | Z=4.32673 |
| | Levant_N | Iran_Hajji_Firuz_ChL | 4.86599e-05 +/- 1.4465e-05 | Z=3.36398 | 137.114 +/- 31.69 | Z=4.32673 |
| | Levant_N | Iran_N | 0.000100338 +/- 2.90258e-05 | Z=3.45684 | 137.114 +/- 31.69 | Z=4.32673 |
| | Levant_N | Iran_Seh_Gabi_ChL | 5.88618e-05 +/- 1.79068e-05 | Z=3.28712 | 137.114 +/- 31.69 | Z=4.32673 |
| | Levant_N | Iran_Tepe_Hissar_ChL | 7.00514e-05 +/- 1.97826e-05 | Z=3.54107 | 137.114 +/- 31.69 | Z=4.32673 |
| | Natufian | Iran_Hajji_Firuz_ChL | 5.70439e-05 +/- 1.39645e-05 | Z=4.08492 | 137.114 +/- 31.69 | Z=4.32673 |
| | Natufian | Iran_Seh_Gabi_ChL | 6.75996e-05 +/- 1.61309e-05 | Z=4.19069 | 137.114 +/- 31.69 | Z=4.32673 |
| | Natufian | Iran_Tepe_Hissar_ChL | 7.20109e-05 +/- 1.74077e-05 | Z=4.13672 | 137.114 +/- 31.69 | Z=4.32673 |
| | Natufian | Levant_ChL | 2.77334e-05 +/- 6.72129e-06 | Z=4.1262 | 137.114 +/- 31.69 | Z=4.32673 |
| **Somali** | Gumuz | Iran_Hajji_Firuz_ChL | 0.000534731 +/- 4.17268e-05 | Z=12.815 | 111.895 +/- 8.47926 | Z=13.1964 |
| | Gumuz | Iran_N | 0.000426485 +/- 3.35371e-05 | Z=12.7168 | 111.895 +/- 8.47926 | Z=13.1964 |
| | Gumuz | Iran_Seh_Gabi_ChL | 0.000519512 +/- 4.13437e-05 | Z=12.5657 | 111.895 +/- 8.47926 | Z=13.1964 |
| | Gumuz | Iran_Tepe_Hissar_ChL | 0.000491898 +/- 3.83357e-05 | Z=12.8313 | 111.895 +/- 8.47926 | Z=13.1964 |
| | Gumuz | Yoruba | 5.37239e-06 +/- 3.52896e-07 | Z=15.2237 | 111.895 +/- 8.47926 | Z=13.1964 |
| | Yoruba | Iran_Hajji_Firuz_ChL | 0.000580886 +/- 4.51896e-05 | Z=12.8544 | 111.895 +/- 8.47926 | Z=13.1964 |
| | Yoruba | Iran_N | 0.000466692 +/- 3.65035e-05 | Z=12.7849 | 111.895 +/- 8.47926 | Z=13.1964 |
| | Yoruba | Iran_Seh_Gabi_ChL | 0.000561414 +/- 4.40781e-05 | Z=12.7368 | 111.895 +/- 8.47926 | Z=13.1964 |
| | Yoruba | Iran_Tepe_Hissar_ChL | 0.000535043 +/- 4.1484e-05 | Z=12.8976 | 111.895 +/- 8.47926 | Z=13.1964 |
| **Amhara** | Gumuz | Iran_Hajji_Firuz_ChL | 0.000562794 +/- 2.65181e-05 | Z=21.223 | 82.1949 +/- 4.3637 | Z=18.8361 |
| | Gumuz | Iran_N | 0.000437318 +/- 2.04666e-05 | Z=21.3674 | 82.1949 +/- 4.3637 | Z=18.8361 |
| | Gumuz | Iran_Seh_Gabi_ChL | 0.000520486 +/- 2.42287e-05 | Z=21.4822 | 82.1949 +/- 4.3637 | Z=18.8361 |
| | Gumuz | Iran_Tepe_Hissar_ChL | 0.000507807 +/- 2.39603e-05 | Z=21.1937 | 82.1949 +/- 4.3637 | Z=18.8361 |
| | Yoruba | Iran_Hajji_Firuz_ChL | 0.000558395 +/- 2.66208e-05 | Z=20.9759 | 82.1949 +/- 4.3637 | Z=18.8361 |
| | Yoruba | Iran_N | 0.000430882 +/- 2.05367e-05 | Z=20.9811 | 82.1949 +/- 4.3637 | Z=18.8361 |
| | Yoruba | Iran_Seh_Gabi_ChL | 0.000518222 +/- 2.45038e-05 | Z=21.1487 | 82.1949 +/- 4.3637 | Z=18.8361 |
| | Yoruba | Iran_Tepe_Hissar_ChL | 0.000499491 +/- 2.4004e-05 | Z=20.8086 | 82.1949 +/- 4.3637 | Z=18.8361 |
| **Oromo*** | Gumuz | Iran_Hajji_Firuz_ChL | 0.000582363 +/- 3.36577e-05 | Z=17.3025 | 101.363 +/- 9.11284 | Z=11.1231 |
| | Gumuz | Iran_N | 0.000467779 +/- 2.62823e-05 | Z=17.7983 | 101.363 +/- 9.11284 | Z=11.1231 |
| | Gumuz | Iran_Seh_Gabi_ChL | 0.000568937 +/- 3.24979e-05 | Z=17.5069 | 101.363 +/- 9.11284 | Z=11.1231 |
| | Gumuz | Iran_Tepe_Hissar_ChL | 0.000529892 +/- 3.00147e-05 | Z=17.6544 | 101.363 +/- 9.11284 | Z=11.1231 |
| | Yoruba | Iran_Hajji_Firuz_ChL | 0.000593155 +/- 3.52004e-05 | Z=16.8508 | 101.363 +/- 9.11284 | Z=11.1231 |
| | Yoruba | Iran_N | 0.000476165 +/- 2.74481e-05 | Z=17.3478 | 101.363 +/- 9.11284 | Z=11.1231 |
| | Yoruba | Iran_Seh_Gabi_ChL | 0.000574846 +/- 3.3418e-05 | Z=17.2017 | 101.363 +/- 9.11284 | Z=11.1231 |
| | Yoruba | Iran_Tepe_Hissar_ChL | 0.000532723 +/- 3.08705e-05 | Z=17.2567 | 101.363 +/- 9.11284 | Z=11.1231 |
| **Wolayta*** | Gumuz | Iran_Hajji_Firuz_ChL | 0.0005344 +/- 3.32157e-05 | Z=16.0888 | 113.486 +/- 11.8732 | Z=9.55815 |
| | Gumuz | Iran_N | 0.000423873 +/- 2.69293e-05 | Z=15.7402 | 113.486 +/- 11.8732 | Z=9.55815 |
| | Gumuz | Iran_Seh_Gabi_ChL | 0.000502396 +/- 3.1138e-05 | Z=16.1345 | 113.486 +/- 11.8732 | Z=9.55815 |
| | Gumuz | Iran_Tepe_Hissar_ChL | 0.000491661 +/- 3.06964e-05 | Z=16.0169 | 113.486 +/- 11.8732 | Z=9.55815 |
| | Yoruba | Iran_Hajji_Firuz_ChL | 0.000558959 +/- 3.50025e-05 | Z=15.9691 | 113.486 +/- 11.8732 | Z=9.55815 |
| | Yoruba | Iran_N | 0.000439468 +/- 2.76294e-05 | Z=15.9058 | 113.486 +/- 11.8732 | Z=9.55815 |
| | Yoruba | Iran_Seh_Gabi_ChL | 0.000522735 +/- 3.19422e-05 | Z=16.365 | 113.486 +/- 11.8732 | Z=9.55815 |
| | Yoruba | Iran_Tepe_Hissar_ChL | 0.000509086 +/- 3.14997e-05 | Z=16.1616 | 113.486 +/- 11.8732 | Z=9.55815 |

*Also have a second more recent admixture event

**Table S3. Testing for admixture in modern populations using ancient sources via Malder.**
Arabian.core represents both Emirati and Saudi core populations combined.

| Test | A | | | | | |
|---|---|---|---|---|---|---|
| | Sidon_BA | Megiddo_MLBA | Jordan_BA | Ashkelon_LBA | Egypt_prePtolemaic | Armenia_EBA |
| Assyrian.HO | 2.94E-20 | 2.07E-104 | 5.31E-53 | 1.36E-15 | 1.18E-30 | 1.64E-04 |
| BedouinA.HO | **2.71E-01** | 1.94E-44 | 1.56E-15 | 3.71E-06 | 1.18E-11 | 2.34E-28 |
| BedouinB.HO | 2.13E-03 | 4.51E-23 | 2.93E-08 | 2.56E-04 | 6.82E-06 | 5.56E-41 |
| Druze.HO | 1.52E-04 | 9.82E-70 | 4.60E-29 | 4.00E-09 | 1.74E-18 | 6.78E-18 |
| Iraqi | 4.43E-15 | 5.51E-77 | 1.20E-38 | 3.27E-14 | 2.14E-27 | 1.31E-11 |
| Jew_Iraqi.HO | 7.62E-11 | 2.80E-59 | 8.75E-35 | 2.29E-11 | 6.65E-24 | 1.45E-07 |
| Jordanian.HO | **2.27E-01** | 9.73E-34 | 3.41E-17 | 4.63E-06 | 1.08E-13 | 6.16E-23 |
| Jordanian | **2.46E-01** | 1.88E-21 | 5.12E-13 | 2.13E-04 | 1.00E-09 | 1.30E-16 |
| Kurd | 3.88E-33 | 1.23E-90 | 5.00E-61 | 2.26E-24 | 6.73E-39 | 5.50E-07 |
| Lebanese_Christian.HO | **7.55E-02** | 1.25E-40 | 9.87E-20 | 6.38E-06 | 1.14E-13 | 2.57E-19 |
| Lebanese_Muslim.HO | 5.87E-04 | 7.63E-58 | 2.26E-26 | 5.34E-08 | 6.21E-18 | 1.39E-15 |
| Omani | **1.90E-01** | 1.21E-14 | 2.53E-11 | 9.64E-04 | 6.66E-12 | 6.84E-11 |
| Palestinian.HO | **7.07E-02** | 2.62E-53 | 2.10E-20 | 1.50E-07 | 3.61E-14 | 2.37E-26 |
| Saudi.core | 1.37E-02 | 5.17E-46 | 1.10E-18 | 6.47E-08 | 1.21E-12 | 2.04E-32 |
| Saudi | **6.06E-02** | 1.05E-22 | 5.74E-09 | 1.87E-04 | 7.16E-08 | 1.26E-35 |
| Saudi.HO | **6.62E-02** | 4.06E-32 | 2.08E-14 | 1.81E-06 | 2.50E-10 | 3.70E-32 |
| Syrian | 4.01E-09 | 2.81E-79 | 4.42E-37 | 1.27E-10 | 3.12E-24 | 1.36E-12 |
| Syrian.HO | 3.42E-03 | 4.63E-43 | 1.99E-22 | 7.82E-08 | 2.91E-17 | 6.81E-13 |
| Emirati.core | **5.48E-02** | 2.11E-38 | 3.53E-17 | 1.53E-07 | 3.59E-14 | 9.13E-29 |
| Emirati | 2.61E-34 | 6.51E-148 | 5.78E-66 | 2.07E-23 | 3.61E-39 | 1.43E-21 |
| Yemeni | 5.10E-03 | 4.67E-27 | 2.24E-10 | 3.69E-05 | 8.40E-08 | 4.19E-40 |
| Yemeni.HO | 9.69E-03 | 1.81E-32 | 2.06E-19 | 1.48E-08 | 1.42E-15 | 6.47E-19 |
| Egyptian.DG | 9.42E-03 | 4.41E-32 | 2.83E-10 | 6.17E-04 | 3.48E-07 | 8.84E-43 |
| Egyptian.HO | 7.40E-03 | 1.07E-27 | 2.50E-09 | 5.95E-04 | 2.43E-07 | 3.85E-42 |
| Amhara.DG | 1.97E-07 | 4.24E-08 | 2.96E-04 | 2.29E-03 | 4.58E-02 | 3.83E-29 |
| Wolayta.DG | 3.31E-03 | 2.59E-03 | **6.79E-02** | 3.17E-02 | **1.03E-01** | 7.76E-14 |
| Somali.DG | 4.64E-07 | 1.44E-05 | 2.52E-03 | 9.62E-04 | **1.19E-01** | 2.24E-20 |
| Somali.HO | 5.73E-05 | 2.30E-04 | 1.24E-02 | 9.02E-03 | **1.74E-01** | 6.05E-17 |
| Oromo.DG | 2.35E-05 | 5.74E-07 | 7.48E-04 | 2.93E-03 | 1.29E-02 | 4.92E-23 |
| Oromo.HO | 2.33E-04 | 1.32E-02 | **6.61E-02** | **7.90E-02** | **5.03E-01** | 5.12E-13 |
| Masai.HO | **1.96E-01** | **6.03E-01** | **6.62E-01** | **3.38E-01** | **5.62E-01** | 5.42E-03 |
| Datog.HO | 1.25E-02 | 2.70E-02 | **5.57E-02** | **9.29E-02** | **2.70E-01** | 6.77E-06 |

**Table S4. Modelling Middle Easterners and East Africans as descending from A, EHG, and Mota.** Showing qpAdm *P* value for rank=2 and testing with different A. Bold values (p-value>0.05) indicate the model cannot be rejected. We tested if we can model our populations as deriving ancestry from one of the sampled regional Bronze Age populations, and found that the Middle Bronze Age population from Sidon (Sidon_BA) could be a source of ancestry for several modern Levantine and Arabian populations (See also Table S5). Our phylogenetic modeling (Figure 3D and S11) suggests that modern Levantines could have directly derived their ancestry from a Sidon_BA-related population; however, the best model for Arabians is an ancestral population distantly related to Sidon_BA harbouring Natufian-like and ancient Iran-related ancestry. We used the following outgroups in the qpAdm test: Ust_Ishim, Kostenki14, Papuan, WHG, CHG, Mbuti, Levant_N and Iran_N.

| | Ancestry proportions | | | Standard Error | | |
|---|---|---|---|---|---|---|
| **Test** | **Sidon_BA** | **EHG** | **Mota** | **Sidon_BA** | **EHG** | **Mota** |
| BedouinA.HO | 0.834 | 0.061 | 0.104 | 0.012 | 0.011 | 0.004 |
| Jordanian.HO | 0.848 | 0.084 | 0.068 | 0.014 | 0.012 | 0.005 |
| Jordanian | 0.895 | 0.08 | 0.025 | 0.019 | 0.016 | 0.006 |
| Lebanese_Christian.HO | 0.91 | 0.088 | 0.003 | 0.015 | 0.013 | 0.005 |
| Omani | 0.85 | 0.043 | 0.107 | 0.02 | 0.017 | 0.009 |
| Palestinian.HO | 0.862 | 0.072 | 0.065 | 0.012 | 0.011 | 0.004 |
| Saudi | 0.809 | 0.036 | 0.155 | 0.013 | 0.011 | 0.005 |
| Saudi.HO | 0.895 | 0.054 | 0.05 | 0.015 | 0.013 | 0.005 |
| Emirati.core | 0.931 | 0.031 | 0.038 | 0.015 | 0.013 | 0.005 |
| Masai.HO | 0.177 | 0.002 | 0.821 | 0.02 | 0.017 | 0.008 |
| | **Egypt_prePtolemaic** | **EHG** | **Mota** | **Egypt_prePtolemaic** | **EHG** | **Mota** |
| Datog.HO | 0.299 | 0.005 | 0.696 | 0.026 | 0.018 | 0.012 |
| Masai.HO | 0.235 | -0.009 | 0.775 | 0.025 | 0.017 | 0.011 |
| Oromo.HO | 0.431 | 0.003 | 0.566 | 0.023 | 0.016 | 0.01 |
| Somali.DG | 0.48 | -0.011 | 0.531 | 0.019 | 0.014 | 0.009 |
| Somali.HO | 0.458 | -0.009 | 0.551 | 0.02 | 0.014 | 0.009 |
| Wolayta.DG | 0.459 | -0.012 | 0.553 | 0.019 | 0.013 | 0.008 |

**Table S5. Ancestry proportions from successful models tested in Table S4.** Showing models involving Sidon_BA or Egypt_prePtolemaic as one of the ancestry sources.
